# Genetic analysis of 7,000 year old preserved goat leather from Cueva de los Murciélagos (Albuñol, Spain)

**DOI:** 10.64898/2025.12.01.691547

**Authors:** Jolijn A. M. Erven, Zoé Robinet Guyet, Louis L’Hôte, Valeria Mattiangeli, Eva Fernández-Domínguez, Geordie Laidlaw, Rafael M. Martínez Sánchez, Blas Ramos Rodríguez, Maria Herrero-Otal, Pedro Henríquez Valido, José Antonio Lozano Rodríguez, The VarGoats consortium, Daniel G. Bradley, Kevin G. Daly, Francisco Martínez-Sevilla

## Abstract

Advances in ancient DNA research have expanded the range of materials from which genetic information can be recovered, enabling the analysis of atypical materials. These often preserve both host and exogenous DNA, for example due to handling or native microbes, offering a window into past diversity. Here we report one such exceptional material: a well-preserved goat leather specimen from Cueva de los Murciélagos (Albuñol, Spain), associated with an Early Neolithic context. Archaeozoological analyses show clear evidence of tanning and anthropogenic modification of the skin, while radiocarbon dating confirms it to be ∼7,250-7,000 years old. We generated genome-wide data from the leather (Murciélagos1) and demonstrated that it derives from a domestic goat of European ancestry. Comparisons with modern goats show highest affinity to the Bermeya breed from northern Spain, demonstrating genetic connectivity in goat herds from the Neolithic to present day Iberia. We additionally recover fragmented DNA from the leather deriving from human and canid sources, the former likely reflecting handling/wearing of the leather and the latter indicating post-depositional disturbance. Murciélagos1 represents one of the oldest nuclear genome data from organic remains outside of high altitude, high latitude regions, and demonstrates the value of ancient DNA analysis in exceptional cases of organic preservation.

## Introduction

The recovery of DNA molecules from ancient and pre-modern biological remains has transformed our understanding of the past. For example, ancient DNA (aDNA) has uncovered previously-unknown relatives of humans (1), located the likely proximal source region for the Black Death in Eurasia (2), and allowed horse husbandry intensification to be quantified by a shortening of generation time (3). Coincidental advances in sequencing technologies have permitted the study of aDNA at increasing scale but typically focus on a small number of material types. These sources are well justified by the taphonomic realities of what tends to be preserved (i.e. inorganic tissue) and in which of those tissues DNA survives best, such as the cochlear part surrounding the osseous inner ear within the *pars petrosa* of the skull temporal bone (4, 5), auditory ossicles (6), and tooth cementum.

However, many studies have also explored atypical materials for their potential to preserve aDNA over millenia. For example, parchment (processed animal skin) has proven to be a source of the animal itself, the humans in physical contact with the manuscript, and the microbes associated with both (7). Mollusc shells (8) and bird feather barbs (9) are just a few examples of the materials that can retain DNA molecules and act as time capsules to directly observe past genetic diversity (10). Mineralized dental plaque (calculus) has also been realized as a source of target DNA and the oral microbiome of ancient peoples (11) and animals (12). These atypical materials allow the recovery of both target DNA and microbiomes, illuminating not only past diversity but also host-microbiome interactions (10).

In exceptional cases, preservation conditions allow organic tissues to remain relatively intact for hundreds or even thousands of years. Indeed, the first reported sequenced ancient DNA was from dried quagga (*Equus quagga quagga*) muscle tissue (13). Environments that can postpone the typical post-mortem degradation of soft tissues include permafrost and anhydrous locations. There are several instances of thousands-of-years-old hair providing endogenous DNA molecules (14–18). Preserved skin has also proven a rich source of ancient biological inference, including genetic data from historic dogs (19, 20), rock pigeons (*Columba livia*) (21), and Egyptian mummies (22, 23); gene expression patterns of the Tasmanian tiger (*Thylacinus cynocephalus*) (24) and the woolly mammoth (*Mammuthus primigenius*) (25); and the genomic architecture of the woolly mammoth (16). Permafrost conditions have also allowed the recovery of aDNA from soft tissue of a Late Pleistocene wolf (26). At lower latitudes, a Neolithic legging of goat skin from the Swiss alps and a naturally-mummified sheep leg from the salt mine of Chehrābād, Iran, provided a 4,200 and 1,600 year old genome, respectively (27, 28).

One such case of exceptional preservation is the Cueva de los Murciélagos site (Albuñol, Granada) in southeastern Iberia (Fig. 1A), where there is outstanding preservation of organic materials ascribed to different phases of Late Prehistory by desiccation (29, 30). The site is a karstic cave in the lower part of the Angosturas Gorge at an altitude of 450 m and 7 km from the actual Mediterranean coast. The shelter of the cave is 15 m wide with an east-facing entrance; the interior part is around 60 m long and 30 m wide, with a 48 m drop.

**Figure 1.**
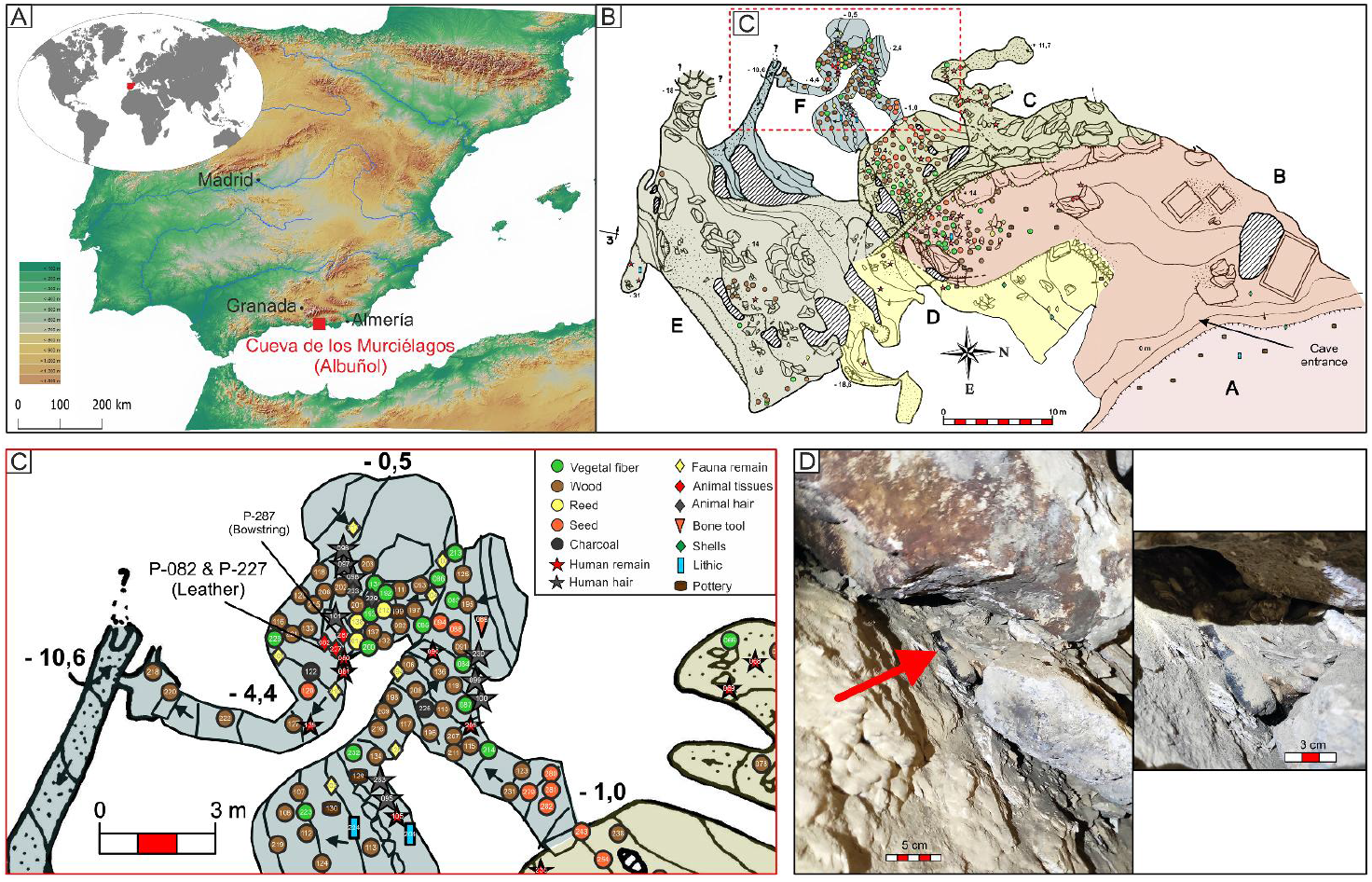
Cueva de los Murciélagos. A) Location of Cueva de los Murciélagos (Albuñol) in the south of the Iberian peninsula; B) Cave plan showing the delimitations used for the survey and for mapping the distribution of archaeological materials from the 2022-2023 surveys (modified from the free access original topography made in 1984 by Gónzalez Rios, M., Menjibar Silva, J.L. and García Ligero, M.; C) Close-up of Sector F showing the location of the tanned skin/leather studied in this paper and the bowstring, both of which were dated; D) Photograph of the leather at the moment of its discovery, embedded in a stone wall built by 19th-century miners.

The first information about Cueva de los Murciélagos dates from 1857, following the interest shown by a mining company in extracting galena from inside the cave. These workers recovered partially mummified human remains, as well as clothing, baskets, and other wooden objects. In addition, a large number of plant artefacts, lithic tools, and ceramics were retrieved, as well as archaeozoological remains (31). Unfortunately, these objects were found disturbed in the cave at the time of their discovery; the majority are now deposited at the Museo Arqueológico Nacional de Madrid (MAN) and the Museo Arqueológico y Etnográfico de Granada (MAEGr). Radiocarbon dating have placed the sequence of the cave funerary use, with a first phase between 9,500-9,000 cal BP (Mesolithic - hunter-gatherer populations); a second phase between 7,200-6,700 cal BP (Early Neolithic - first farming populations); a third phase between 6,700-6,000 cal BP (Middle Neolithic); and a fourth phase between 6,000-5,700 cal BP (Late Neolithic). Finally, there is a Bronze Age period between 3,900-3,700 cal BP. The Cueva de los Murciélagos site has been interpreted as a burial place in those periods, where goods placed likely have a symbolic significance (30).

The fieldwork in the cave, in the framework of the MUTERMUR project, was carried out as an intensive survey, and allowed for the recovery of different archaeological materials in the miners’ waste dumps (Fig. 1B). During the surface survey, two fragments of animal skin (CM22/SF/P-082 and CM22/SF/P-227) likely belonging to the same individual were found together beneath a block moved by miners in a deep area of the cave, defined as Sector F (Fig. 1 C & D). Initial archaeozoological assessment of the skin ascribed it to the genus *Capra*.

Accordingly, the objective of this study is to evaluate the biological and chronological identity of this skin object from Cueva de los Murciélagos, named Murciélagos1, to determine whether it is a Neolithic representative of the domesticated species *Capra hircus* or the wild species *Capra pyrenaica*. To answer this question, we combine population genetic analyses, aimed at providing biomolecular evidence for the specimen’s taxonomic classification, with radiocarbon dating, to situate the specimen within its archaeological and temporal context. Through this integrated approach, we aim to shed light on the exploitation and management of goats during the Neolithic in Iberia.

## Results

The materials analyzed here correspond to two different fragments of the same animal skin (Table S1). Fragment P-082 is an irregularly and asymmetrically tanned hide - from this point referred to as leather - tending towards a rectangle measuring 346 mm long by 202 mm wide. The second fragment P-227 is a piece of 226 mm long by 138 mm. The leather retains hair on one surface, while the opposite side displays traces consistent with tanning treatment as well as four perforations showing clear evidence of anthropogenic modification (Fig. 2). These perforations have been identified as anthropogenic, based on the distinction that breaks produced by insect activity exhibit irregular, stepped edges typical of gradual biological degradation, while anthropogenic perforations show smooth, sharply defined margins consistent with deliberate human modification (Fig. 2C & 2D).

**Figure 2.**
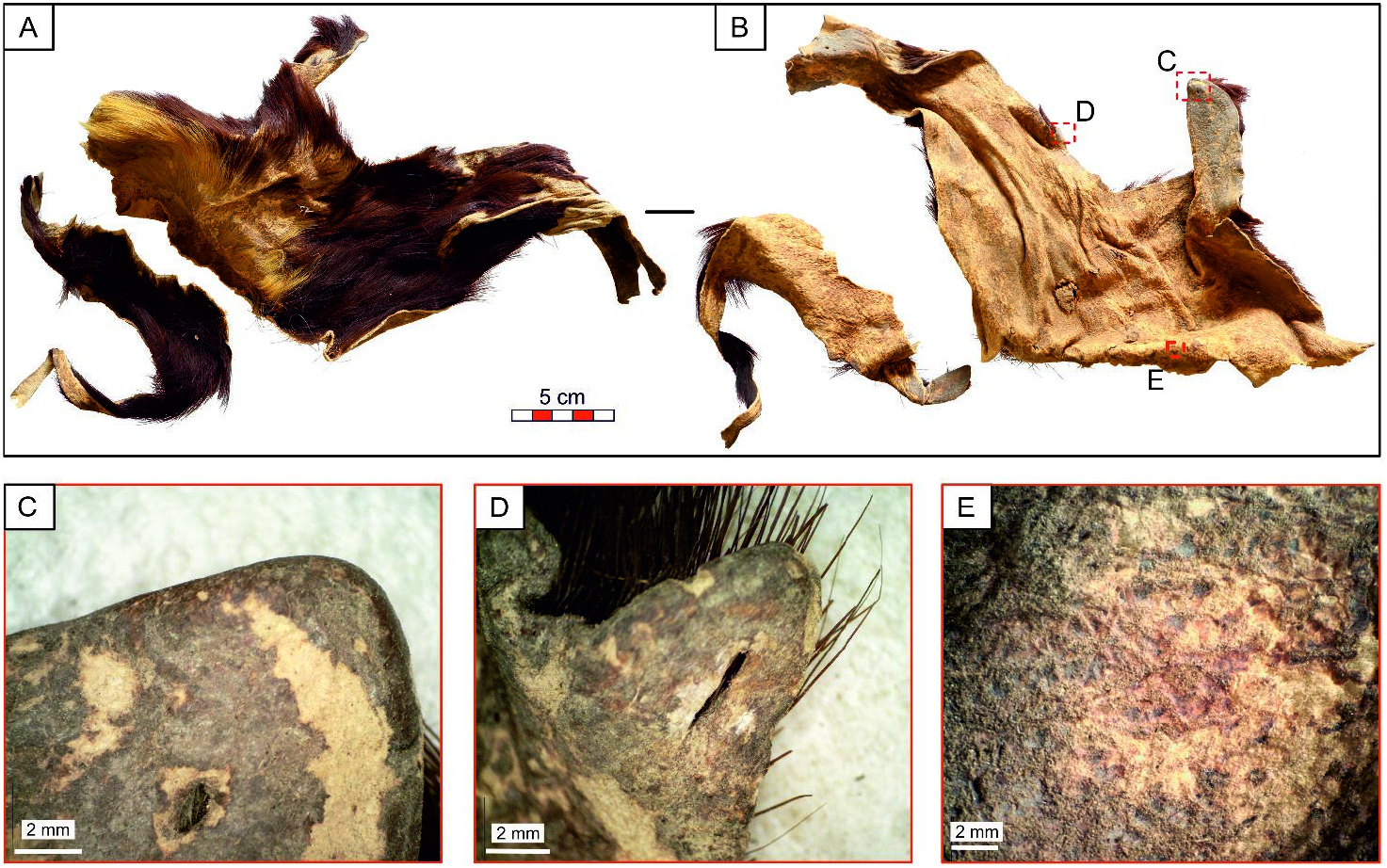
Goat leather from Cueva de los Murciélagos of Albuñol. A) hair side of leather; B) flesh side of leather; C & D) Example of anthropic perforation and tanned edges; E) Tanned surface with traces of softening from use.

The radiocarbon dating of the Cueva de los Murciélagos leather (Ua-78252) - from this point referred to as Murciélagos1 - yielded an age of 7,246-7,008 cal BP (2σ calibrated range; Fig. 3A). In the context of the cave’s chronological framework, this estimated age places this object within the earliest phase of the Early Neolithic (Fig. S1, Table S2). This phase of material deposition has been identified in other objects, such as bowstrings (30). Together with the presence of human remains, this suggests that the area where the leather was found was used as a burial site during the Early Neolithic, and that these objects were deposited as grave goods.

**Figure 3.**
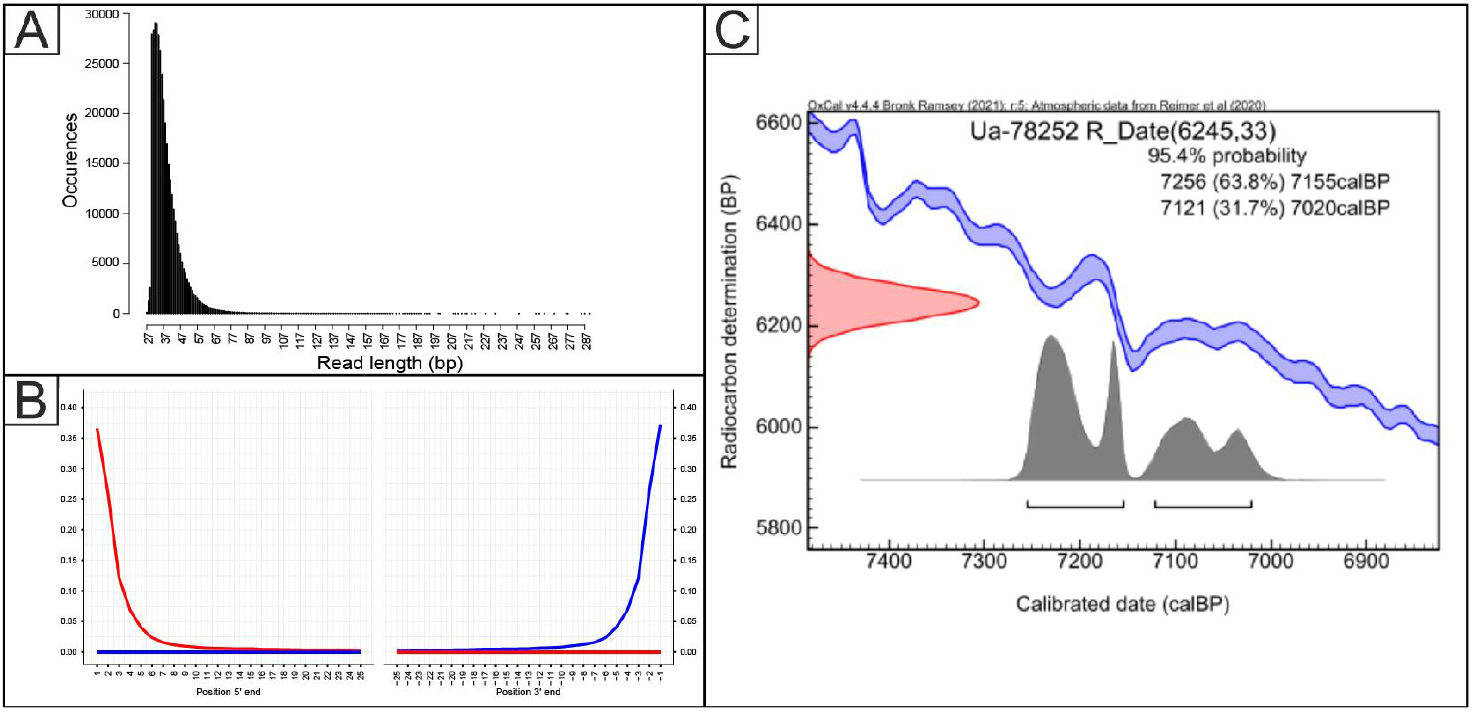
Temporal assessment of Murciélagos1. A) Read length distribution; B) aDNA damage, C>T (red) and G>A (blue) frequency of misincorporation rates at the 3’ and 5’ end of reads; C) Radiocarbon date calibrated in OxCal 4.4 software(32) using the IntCal20 terrestrial calibration curve (33).

To determine if Murciélagos1 was domestic *Capra hircus*, wild *Capra pyrenaica* or something else, we aligned sequencing reads across a set of reference genomes using FastQ Screen (34) (Table S3). Initial assessment indicated that most reads were assigned to the goat ARS1 genome (433,227 reads, 46% of total mapped reads, Table S1). However, we also detected some reads aligning to either human and fox reference genomes (Table S1,4). To account for this we removed reads aligned to the goat genome which also aligned to either the fox or human genomes (161,320 and 344,827 reads, contributing 17% and 37% of total mapped reads respectively, Fig. S2-6), leaving 377,825 goat-aligned, unique reads (40% of total mapped reads) after other quality control steps. In sum, we retrieved both mitochondrial DNA (36.68×) and autosomal DNA (0.0049×) from the preserved leather. Aligned reads showed a short average length (37 bp) and substantial postmortem damage patterns compatible with aDNA (Fig. 3, S3-7). We determined the molecular sex of Murciélagos1 to be female (Supplementary Note 1, Fig. S7-8).

We additionally characterized the metagenomic content (see Supplementary Methods) of the non-mammalian reads: the majority fall within the Eukaryota domain (90.4%), and more specifically within the kingdom Fungi (88.1%), with a predominance of the phylum Ascomycota (82.1%; Fig. S9), which is one of the most dominant Fungi in caves (35). We additionally assessed human-assigned reads: after merging all human-aligned reads and genotyping with nf-core/EAGER ((36), Supplementary Note 2, Table S4), only 7,261 SNPs overlapped with the 1,352K SNPs in the Ancient Human DNA Target Enrichment panel (Twist Bioscience), while only 133 reads aligned against human mtDNA (Table S4). Molecular sex of the human DNA was assigned as male; however, due to the low number of reads this assignment should be considered tentative (Fig. S10). Due to insufficient coverage of potentially informative SNPs and lack of phylogenetic resolution, no further analyses were conducted with the human reads.

To probe if the genetic ancestry of Murciélagos1 was consistent with that of a Neolithic Iberian domestic goat, we computed a projection PCA using LASER 2.0 (37). Using the VarGoats modern genome dataset as a background reference (Table S5, (38)), Murciélagos1 falls within European ancestry (Fig. 4), close to modern southern European breeds (Swiss, Italian and French), and overlapping with the modern Swiss breed cluster. In contrast, modern day Spanish breeds are shifted away from the European cluster towards African breeds. From these, the available northern Spanish breed (Bermeya) is closer to the European cluster, while the Palmera breed from the Canary Islands falls within the African cluster (Fig. S11). This genetic affinity in the Spanish goat from the Balearic islands and Catalonia with African goat populations is likely due to African introgression into these Spanish breeds, previously shown to have occurred (39, 40). To further test the domestic status of Murciélagos1, a *D* statistic test (*D*(Out, Neolithic Goat; *Capra pyrenaica*, Modern breeds)) was performed, showing that Murciélagos1 has a closer genetic affinity to modern domestic breeds than with wild *Capra pyrenaica* (Iberian ibex; Fig. S12, Table S6; *D* statistic range 0.068-0.080, Z values 34-234), consistent with Murciélagos1 representing a domestic goat.

**Figure 4.**
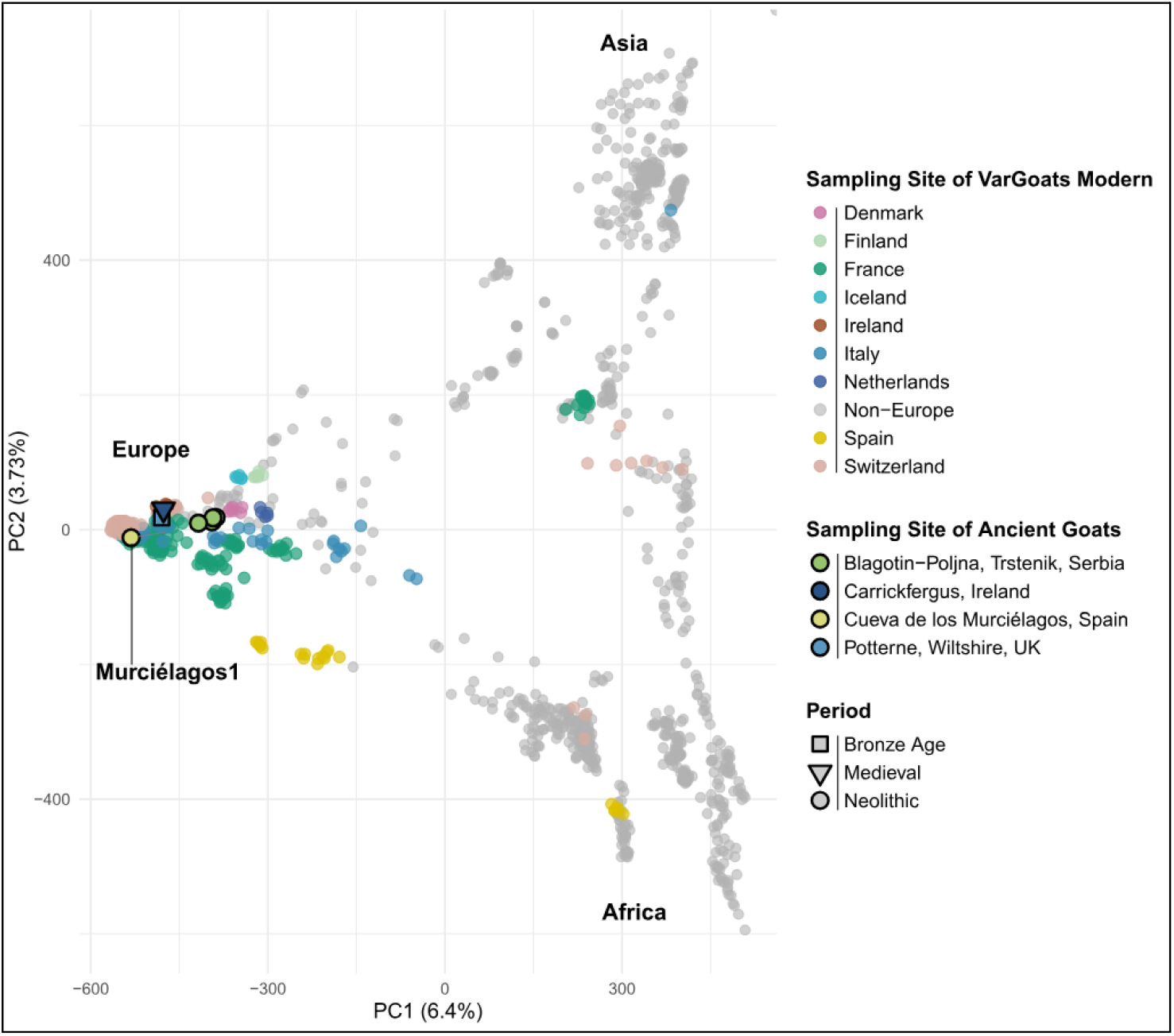
Principal component analysis of the VarGoats Modern dataset and ancient European goats, including our study sample Murciélagos1 goat leather. Each point represents the genetic information of a goat sample. Axes PC1 (6.4%) and PC2 (3.73%) represent the first two principal components, which explain the genetic variation. Grey points correspond to modern non-European samples; those located at the bottom right of the PCA form the African cluster, and those at the top right form the Asian cluster.

Given the low total number of aligned reads, we sought to further confirm the relationship between Murciélagos1 and modern goat breeds. To do this we performed an outgroup *f*_*3*_ analysis to measure the amount of shared genetic drift between Murciélagos1 and modern goat breeds.Consistent with PCA results, Murciélagos1 shows more genetic affinity with European breeds compared to African and Asian breeds (Fig. 5). Notably, the Bermeya breed (BEY) from northern Spain shows the highest shared genetic drift with Murciélagos1 (Table S7). The Valais Copper Neck (Swiss) and Poitevine (French) breeds also show high genetic affinity with Murciélagos1. In summary, these results suggest a degree of population connectivity between the Murciélagos goat and both Spanish and European goat breeds today.

**Figure 5.**
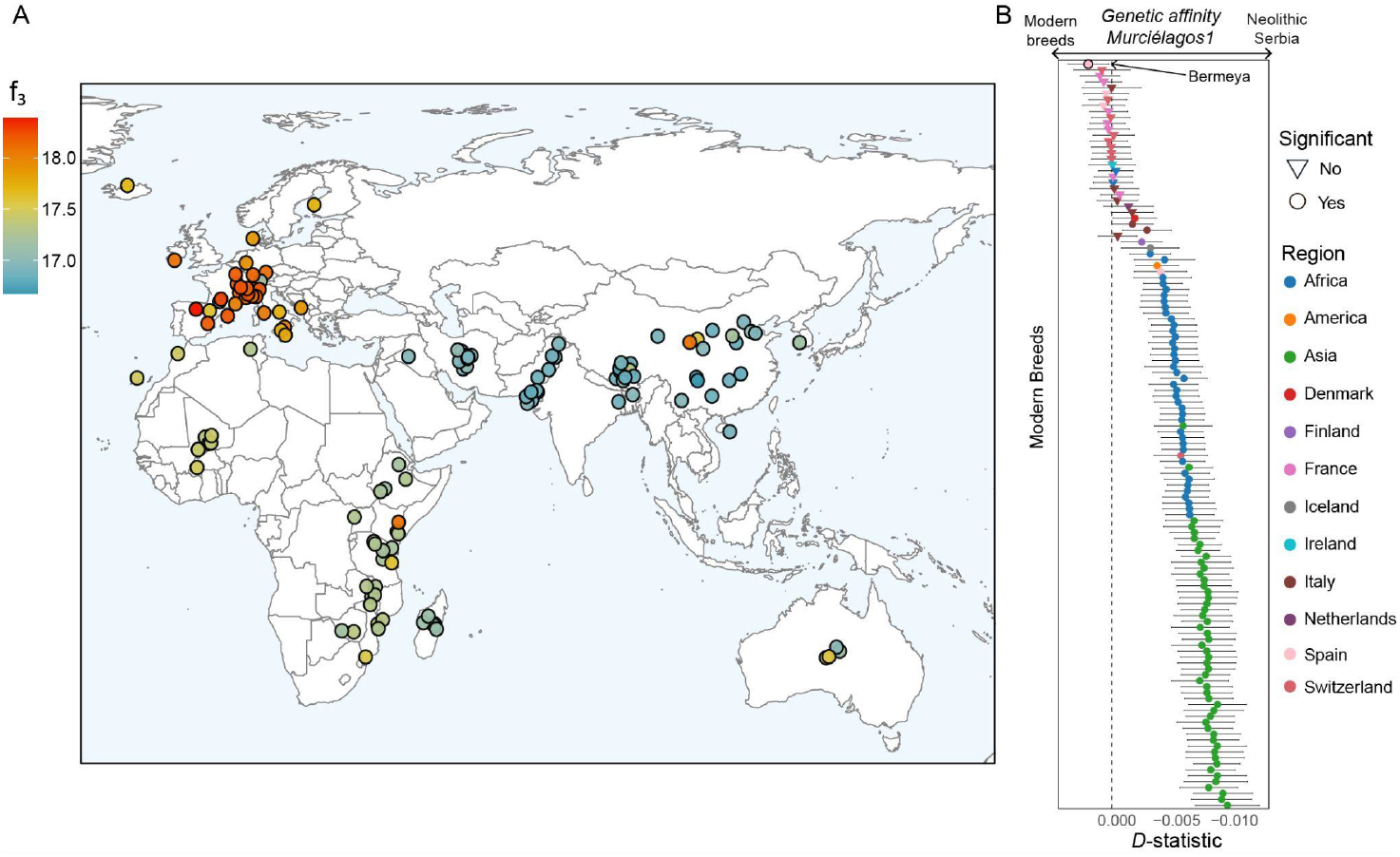
Outgroup f_3_ and D statistics for Murciélagos1. A) Geographical distribution of shared genetic drift (outgroup f_3_) between the Neolithic Murciélagos1 goat from Iberia, and modern domestic goat breeds. The range of the outgroup f_3_ statistic in modern domestic comparisons ranges from 15 to 30. B) D statistics of the form D(Out, Murciélagos1: Neolithic Serbia (Blagotin), Modern breeds). This D test measures affinity of Murciélagos1 with Neolithic Serbia or modern breeds. Negative indicates closer affinity (shared derived alleles) of Murciélagos1 to Blagotin and positive indicates closer affinity towards the modern breed in question (Table S8).

We then used *D* statistics to confirm the close genetic affinity between Murciélagos1 and the northern Spanish Bermeya breed. We tested if ancient individuals shared more genetic affinity with the Bermeya breed compared to other modern breeds (*D*(Sheep, Neolithic Goat; Bermeya, Modern breeds), using Murciélagos1, Neolithic Serbian genomes (∼7936-8190 BP (41)), and a Neolithic East Iranian genome (∼8,055 BP (41)) as comparisons. Murciélagos1 shows greater genetic similarity to the Spanish Bermeya breed than to other breeds today (Fig. S13, Table S8); *D* statistics ranged from −0.00081 to −0.012 (Z value ranged from −1.14 to −18, 5 breeds did not reach significance at |Z| > 3). However, this trend is not significant for some southern European breeds, which also displayed strong similarity with Murciélagos1 in the outgroup *f*_*3*_ analysis. In contrast, a Neolithic Serbian goat shows greater genetic similarity to southern and central European breeds rather than to Bermeya goats, while a Neolithic Iranian genome shows low genetic affinity to modern European breeds.

We additionally tested the relative sharing of derived alleles between Murciélagos1 and either the closest available Neolithic goats (Blagotin-Poljna, Serbia) versus the modern breeds with the *D* statistics test *D*(Out, Murciélagos1; Neolithic Serbia, Modern breeds), then ranked results based on degree of greater allele sharing with Murciélagos1. The Bermeya breed shows highest relative allele sharing with Murciélagos1 among modern breed comparisons (Fig. 5B, Table S9, *D*=0.0023, Z=4). Other southern European breeds show a similar pattern of relatively high allele sharing with Murciélagos1, although their test statistics do not reach significance likely due to the low sequencing depth available for Murciélagos1 (*D* statistic range 0.0015-0.000654, Z score 2.52-1.12). Combined these analyses demonstrate that Murciélagos1, unlike other published ancient goats, shows a high genetic affinity with Spanish breeds, particularly the Northern Spanish Bermeya breed, suggesting that Murciélagos1 belonged to or was related to a population that gave rise to or with a direct genetic connection to Spanish goat breeds today. A final *D* statistic of the form *D*(Out, Neolithic; Moroccan, European/African Modern breeds) demonstrated that Murciélagos1 shows no genetic affinity towards African breeds (Fig. S14, Table S10).

Finally, we assessed the mitochondrial genome of Murciélagos1, placing it within the known diversity of modern and other ancient goat samples. Murciélagos1 belongs to haplogroup A (Fig. S15), the haplogroup most commonly observed in European goats in the Neolithic today (41). The bootstrap value at the haplogroup A node is 95, which provides strong support for this assignment. However, the precise position of Murciélagos1 within this haplogroup is not well resolved, with low bootstrap values, indicating uncertainty regarding its relationship to other sequences within haplogroup A.

## Discussion

Murciélagos1 (∼7,250-7,000 years old) represents the to-date oldest nuclear genomic data reported from dessicated preserved skin remains outside of permafrost or frozen context. Goat DNA has been recovered from a 4,200 year old leather legging (28). mtDNA data has been reported from 4,000-3,500 year old human mummified remains from the Tarim Basin in China, but from dental and skeletal parts (42). The leather clothing of the Tyrolean Iceman (∼5,300 BP) has yielded mtDNA, but these derive from a glacial context (43). Similarly, mtDNA amplification has been reported of Neolithic leggings made of goat skin, dating to ∼4,700BP, but were found in high altitude context (28). DNA has also been recovered from the dessicated soft and hard tissues of Egyptian mummies, but these are comparatively recent (∼3,400-1,500 BP) (23), as are reports of genetic data from a variety of tissues from Egyptian ibis (*Threskiornis aethiopicus) (44)*. Additionally, 4,000 year old DNA has been reported from preserved human hair, from Greenland (45) and Sudan (15). Finally, genomic data has been obtained from a naturally mummified sheep leg, but dates to ∼1,600 BP (27). In contrast, Murciélagos1 (∼7,200 years BP) represents skin tissue preserved through desiccation in a low latitude, cave environment, which has retained sufficient genetic data for nuclear genome exploration.

While study of the materials is on-going, an initial assessment of the faunal bone assemblage from Cueva de los Murciélagos sheds light on Neolithic strategies for selecting and processing animal raw materials. There was marked preference for long and flat bones from medium and large ruminants - particularly domestic goats, but also bovids and red deer (46). Goat metapodia and tibiae were frequently chosen to craft pointed implements, spatulas, and toothed tools, using techniques such as percussion, splitting, abrasion, and longitudinal grooving. Traces of intensive use and microscopic wear patterns underscore both the functional efficiency of goat bones and the advanced craftsmanship of the prehistoric makers (46). Additionally, bowstrings composed of animal sinews and preserved at Cueva de los Murciélagos were identified by ZooMS as *Capra* species (30). Radiocarbon dating of a cord (Beta-684048) showed a date between 7,239-6,999 cal BP, contemporary with Murciélagos1. Our genetic confirmation that Murciélagos1 is a domestic goat provides evidence that, at least for this specimen, domestic flocks were exploited in the development of this material culture, rather than Iberian ibex (*Capra pyrenica*) indigenous to the Peninsula prior to the arrival of husbandry.

The introduction of managed animals to Iberia marked a pivotal shift in subsistence strategies, reflecting the spread of Neolithic husbandry and human-animal interactions across the region. Within a few centuries, livestock herds came to constitute the majority percentage of zooarchaeological assemblages in the Mediterranean and southern Iberia during the Early Neolithic (47–49). It does not seem possible to draw reliable patterns by considering the Peninsula as a whole (50). In fact, livestock herding does not seem to play a major role in assemblages from the northern and northwestern Peninsula until the following millennium (7th millenium BP), when the importance of domestic herds approaches that observed in other Iberian regions (49, 51). From the Early Neolithic onwards, coastal sites continue to show faunal spectra inherited from hunter-gatherer economies, to which the domestic component was gradually incorporated, as seen at Cova de les Cendres (Alacant, eastern Iberia) (52), Nerja (Vestíbulo and Torca sites) (53), and Zacatín (Gualchos-Castell de Ferro) (54) in the southern Iberian Peninsula.

The genetic data recovered from Murciélagos1 offers the first glimpse at the genetic makeup of goat populations from Neolithic southern Iberia. This piece shows genetic ancestry consistent with European domestic goat populations, in agreement with the archaeological evidence of the use of managed animals and crops at the site (29, 55). Murciélagos1 has the highest affinity with modern southern European breeds, particularly with the northern Spanish Bermeya breed. This implies that Murciélagos1 was part of, or closely related to, a population that was at least partially ancestral to the Bermeya breed. Yet, Murciélagos1 does not show a high affinity with southern Spanish breeds, likely a consequence of later African introgression in these breeds (39, 40). While there is African ancestry in Bermeya goats, it is notably lower compared to the other southern Spanish breeds (39, 40).

The presence of African ancestry in Spanish goat breeds is evident, yet the question on when this happened remains unclear. There is archaeological and paleogenomic evidence of connections between the southern Iberian Peninsula and the northwest coast of Africa (56–59), and one study indicates a synchronous emergence of the Neolithic in southern Iberia and northwestern Africa (57). Earliest evidence of domestic goats in southern Iberia is dated to ∼7,500 B P (60), ∼7,500 BP in Tunisia (61), and ∼7400-7000 BP in Algeria (62) and Morocco (58, 63, 64). The arrival and dispersal routes of these Neolithic communities have been widely debated (61, 65), with proposed scenarios ranging from a dual-dispersal model to either an Iberian (58, 66) or an African diffusion (56). Our data suggests that African goat introgression was not present in the Early Neolithic in the region around Cueva de los Murciélagos; Murciélagos1 does not show genetic affinity with African breeds compared to European breeds (Fig. S14). However, we note that Murciélagos1 has low sequencing depth, and additional data may refine our understanding. The earliest evidence of African human gene-flow in Iberia, around 4,500 years BP, comes from Camino de las Yeseras in central Iberia, with a human male carrying African ancestry (67), contemporaneously ivory and ostrich eggshells appear in Iberian tombs further elucidating African influence in Iberia during this period (58). This, together with the absence of African goat affinity in Murciélagos1 (Fig. S13-14) may suggest that there was no Africa-to-Iberia migration (or movement of goats) in the Early Neolithic of Iberia, and that the population ancestral to Murciélagos1 had a European rather than African origin. Despite this absence of African ancestry in both Iberian humans and goats, Neolithic European farmer ancestry was detected across the Strait of Gibraltar around 7,400 BP (58, 68), suggesting a unidirectional connection from Iberia to the Tingitan Peninsula and possibly the establishment of a founding population in this region. More genetic data from livestock remains in Iberia and northwestern Africa may further contribute to resolving the debate surrounding the Neolithic dispersal routes and the contact between Iberia and northwestern Africa.

In addition to the host organism (goat), the occurrence of human and fox-aligning reads (Table S1) provides genetic evidence that the Murciélagos1 leather may have been handled and disturbed post-mortem. Given the absence of directly associated skeletal remains and the presence of mammal modifications (Fig. 2), the leather was likely modified post-mortem by humans. While analysis of human remains are on-going, the leather was likely part of a funerary assemblage. There are many human remains within the same context (Fig. 1C); the tanned nature of the leather suggests it was probably worn by one or multiple individuals, and eventually deposited as part of a person’s funerary assemblage. The handling of the goat leather resulted in exogenous contamination of human DNA, with evidence of ancient authenticity of the recovered DNA (Fig. S3-6). The presence of fox (or possibly canid) aligning reads is a curiosity; while there is evidence of presence of fox around Cueva de los Murciélagos; their occurrence is low during the Early Neolithic and became more common from the end of the Neolithic till the end of the Bronze Age (∼5,000-3,000 years BP) (69). The presence of fox DNA could reflect co-mingling of the leather with the remains of foxes or other canids, or more likely by the disturbance of the funerary context by wild foxes/canids during or after the Early Neolithic. The post-mortem damage and fragmentation of fox-aligned reads in Murciélagos1 (Fig. S6) are reduced compared to goat or human-aligned reads (Fig. S2-4), suggestive of a more recent origin.

In summary, we reported the first nuclear genetic data from domestic goats in Iberia, obtained from a tanned goat leather specimen dated to the late 8th millennium BP. This represents one of the oldest genome data recovered from organic remains outside of high altitude, high latitude conditions. The remarkable preservation of the specimen includes several deliberate perforations on the edges of the leather, perhaps made to allow for wearing of the leather as a garment. While the recovered data is limited, it allows to infer a post-Early Neolithic incursion of African goat ancestry to Spanish goats. Denser genetic sampling both temporally and geographically of ancient caprine remains within Iberia will refine our understanding of this process which shaped goat breeds alive today.

## Materials and Methods

An in-depth description of the materials and methods utilised are provided in the SI appendix, a brief overview is provided here. The material, a tanned goat leather specimen, was consistently handled with powder-free gloves to prevent contamination. A portable digital microscope was used to distinguish between anthropogenic perforations and damage from post-depositional processes. Palaeogenetic work was performed in a dedicated ancient DNA facility in Trinity College Dublin. Researchers minimized potential contamination of the sample using personal protective equipment. DNA was extracted from the leather specimen (Fig. S16); Three skin aliquots (identified as S1-S3) were subjected to a diluted ethanol wash and proteinase digestion. Two extraction approaches were performed for the three aliquots (S1-S3); Tubes S1 and S3 were subjected to a phenol-chloroform extraction (70), labelled PEX. Tube S2 was subjected to a purification protocol using High Pure Viral Nucleic Acid Large Volume silica-based spin columns (71), labelled MEX. Tubes S1-2 were USER treated (72), S3 was not USER treated; this resulted in three different aliquots 1) Partial UDG treatment PEX, 2) partial UDG treatment MEX, and 3) non-UDG treatment PEX. Double-stranded DNA (dsDNA) libraries were then constructed (30). Sequencing was performed on an Illumina NovaSeq 6000 (2 × 100 bp).

Quality control was performed with (73, 74), trimming and collapsing of the reads was performed with AdapterRemoval v2.3.1 (75). Taxonomic composition and contamination were assessed using FastQ Screen v0.15.2 (34), which identified goat, human and fox-associated reads. Collapsed reads were aligned to the goat reference genome ARS1 (76), human reference genome hg39, and fox reference genome (vulvul2-2) with BWA aln (77), using parameters tailored for ancient DNA (-l 1024 -n 0.01 -o 2) (39). Reads aligning to either human or fox were removed from the goat ARS1 bam file (Table S1). Reads aligning to human and goat were removed from the fox bam file, and reads aligning to fox and goat were removed from the human bam file (Table S4). MapDamage (78) was applied to assess aDNA damage patterns (Fig. 2, S2).

A projection Principal Component Analysis (PCA) was performed using LASER v2.4 (37). The PCA reference space and projection transformation were constructed using the VarGoats dataset (43, 44), filtered to retain transversions with a MAF of 5%, resulting in a total of 2,420,382 SNPs, as in (79). Transversion pseudohaploid genotypes were called using ANGSD v0.941-11-g7a5e0db (80), with the -doHaploCall function, restricting the analysis to 8,026,660 autosomal transversion SNPs from the VarGoats dataset. Genotypes were converted with haploToPlink, then merged with the VarGoats dataset, and converted to EIGENSTRAT using ADMIXTOOLS’ convertf tool (81). These files were used to compute outgroup *f*_*3*_ statistics (82) and *D* statistics with ADMIXTOOLS version 7.0.2 (81). Granular breed groupings were used, with domestic sheep as the outgroup. For mitochondrial DNA, collapsed reads were aligned using bwa aln to a circularized version of the goat mitochondrial genome (NC_005044.2), followed by a realignment to a circularized haplogroup representative; consensus sequences were generated with ANGSD v0.941-11-g7a5e0db. Mitochondrial sequences were aligned using MAFFT v7 (83). Mitochondrial trees were built with RAx mL v8.2.12 (84) under the GTR+Gamma substitution model.

To characterize the microbiome, host reads were removed by aligning sequences with Bowtie2 v2.4.4 (85) against a concatenated reference of human and several animal genomes. Unmapped reads were analyzed with KrakenUniq v1.0.4 (86) for metagenomic profiling using a custom database (87). Radiocarbon dating was performed at the Tandem Laboratory, Uppsala University. Lipids were extracted, and the insoluble fraction was isolated through acid and base treatments, then dried and combusted for ^14^C analysis. The resulting date (Ua-78252) was calibrated in OxCal v4.4.4 using the IntCal20 curve (32, 33).

## Acknowledgements

This research has been carried out within the framework of different research projects: ‘De los museos al territorio: actualizando el estudio de la Cueva de los Murciélagos of Albuñol (Granada)’ (MUTERMUR) (CM/JIN/2021-009) and ‘La Cueva de los Murciélagos of Albuñol una ventana única a la Prehistoria del sur de Europa’ (MUTERMUR 2.0) (CM/DEMG/2024-046) financed by the Comunidad de Madrid and Universidad de Alcalá (directed by F.M.S.). Fieldwork at the site was conducted with permission from Delegación Territorial de la Junta de Andalucía de Cultura y Patrimonio Histórico de Granada (EXP: BC.03.143/22 14100 and EXP: BC.03.136/23 15490). We would like to thank all the volunteers and students involved in the fieldwork, and especially technician Sergio Fernández Martín for his supervision and willingness during the work. P.H.V is supported by a postdoctoral contract Juan de la Cierva by the Ministerio de Ciencia, Innovación y Universidades y Agencia Estatal de Investigación (AEI) (JDC2023-051629) and RMMSś work is part of the project “First Settlements in Inland/Central Andalusia” (ERSAND) [Ref.PID2023-152309NA-I00 /funded by the Ministry of Science, Innovation and Universities of the Government of Spain]. This publication has emanated from research conducted with the financial support of Taighde Éireann – Research Ireland under Grant number 21/PATH-S/9515(T); and was supported by the European Research Council under the European Union’s Horizon 2020 research and innovation program (grant no. 885729-AncestralWeave).

## Data availability

The VarGoats genotypes are under embargo until 31st December 2025, and will then be available at ENA under Project accession PRJEB90141. The genetic data generated here is deposited in the European Nucleotide Archive (ENA; accession no. PRJEB104374). Previously published data was used for this work (41, 88, 89).

