## Supplementary_methods-Note_Figures_S1-16 for "Genetic analysis of 7,000 year old preserved goat leather from Cueva de los Murciélagos (Albuñol, Spain)"

### Supplementary Information

**Materials and Methods**

Methods

Material description and digital register.

The material was handled with the utmost care throughout the process of collection and sampling. Powder-free gloves were consistently used to prevent contamination, and the number of individuals involved in its manipulation was kept to a minimum. These measures ensured controlled conditions and reduced the risk of external interference. The fragments were photographed with a Canon 77D camera equipped with a Canon EF-S 18-55 mm lens. More detailed images were captured using a portable digital microscope (Dino-Lite Edge Digital Microscope AM7915MZT) with a magnification range of ×20-220. This approach allowed us to distinguish anthropogenic perforations from damage caused by post-depositional processes and insect activity (Fig. 2).

Sample preparation, DNA extraction and library construction

All palaeogenetic work was performed in a dedicated ancient DNA facility in Trinity College Dublin. Researchers minimized potential contamination of the sample using personal protective equipment (gloves, facemasks, and full-body boilersuits) and frequent cleaning of the environment using diluted sodium hypochlorite solution (0.5%).

A small cutting of specimen P-082 was prepared for DNA extraction by first scraping hair from the skin tissue using a scalpel blade (Fig. S16). The skin was then fragmented by repeated manually cutting using the blade and tearing the skin using metal forceps. The skin was then divided into three fractions (KD413S1-3) and then transferred to separate 2 mL tubes using the scalpel.

Three skin aliquots (tubes S1, S2, S3) each weighing ~50mg were subject to a $H_2O$-ethanol-$H_2O$ wash. One extraction control was included for this and following steps. 1 mL of pre-UVed $H_2O$ was added to each tube, the vortexed, then briefly spun to maximum speed (17,000 g) on a bench centrifuge, and the supernatant removed. This was repeated using 600 µL of 100% laboratory-grade ethanol, and then a final 1 mL $H_2O$ wash. The specimens were subject to one additional spin to maximum speed (17,000 g) and any remaining supernatant removed using a pipette.

5 mL extraction buffer was then prepared from: 4,021 µL $H_2O$, 138 µL NaCl (4 M), 275 µL Tris HCl (1 M), 11 µL EDTA (0.5 M), 275 µL SDS (10%), 30 µL $CaCl_2$ (0.5 M), and 250 µL 1M DTT; the buffer was vortexed briefly to ensure mixing. 450 µL of extraction buffer was added to each sample tube, followed by 50 µL proteinase K. Tubes were parafilmed, briefly vortexed and then incubated at 56°C overnight under constant rotation.

We then applied two extraction approaches for each of the three aliquots (S1-S3). Tubes S1 and S3 were subject to phenol-chloroform extraction (1) and here labelled "PEX". 1 mL phenol was added to each sample tube, the tube vortexed, and then rotated at room temperature for 15 minutes. 1 mL of a 24:1 chloroform-isoamyl solution was then added to fresh, labelled tubes. The aqueous layer of the sample tubes (~1 mL) was carefully removed using a pipette and added to the chloroform-isoamyl solution-containing tubes. Each tube was vortexed and rotated at room temperature for 10 minutes.

The tubes were then centrifuged at maximum speed (17,000 g) for 5 minutes. In parallel, 50 µL NaAce (3 M) was added to fresh, labelled tubes. The aqueous layer of each sample tube (~1 mL) was then transferred to the new tubes. 900 µL of cold (refrigerated) 100% ethanol was then added to each tube, and the tubes left in a refrigerator overnight. The tubes were then spun at maximum speed (17,000 g) for 10 minutes. The supernatant was removed using a pipette and discarded. Tubes were then left on clean paper towels open side down for 60 minutes to evaporate remaining ethanol. Finally, 50 µL EBT (elution buffer + 7.5 µL tween) was added to each tube and mixed thoroughly to resuspend DNA.

For the sample aliquot S2, we followed an established purification protocol using High Pure Viral Nucleic Acid Large Volume silica-based spin columns (Roche), detailed under "Extraction Day 2" at (2) and here labelled "MEX". Briefly, following overnight extraction the sample tube was spun at maximum speed (17,000 g) for 10 minutes, and the supernatant carefully transferred to 50 mL Roche High Pure Extender Assembly spin columns containing 13 mL modified PB (420 µL Sodium Acetate (3 M) and 330 µL NaCl (3 M)). Tubes were spun in a centrifuge with a swing bucket rotor for 2 minutes at 700 g, the tubes rotated, and a final spin for 2 minutes at 600 g. Purification then continued according to the standard protocol, modified so the centrifuge spin speed was reduced 3000 g to all but the final EBT elutions; and for the elutions, two successive 25 µL EBT (heated to 65 °C) elutions were performed, to maximize DNA recovery.

Double-stranded DNA (dsDNA) libraries were then constructed for each sample following (3), using 16.25 µL extracted DNA as detailed previously (4). For extract S3, DNA was not treated with Uracil DNA-glycosylase prior to library construction; all other libraries were treated with 5 µL Uracil DNA-glycosylase for 1 hour at 37°C before library construction. This produced three libraries: one from phenol-chloroform extraction, partial-UDG treatment (S1-PEX1-UDG1), one from the modified EB-based extraction, partial-UDG treatment (S2-MEX1-UDG1), and a third by the phenol-chloroform extraction, non-UDG treatment (S3-PEX1-NUDG1). The resulting libraries were subject to PCR-amplification using unique dual indexes; amplified libraries were pooled and sequenced on an Illumina NovaSeq 6000 (TrinSeq, Dublin).

**Data handling**

Initial quality control of raw FASTQ files was performed with FastQC v0.11.9 (5). FastQC reports were then aggregated with MultiQC v1.27.1 (6) to identify quality issues). Sequence cleaning was performed with AdapterRemoval v2.3.1 (7), removing adapters and low-quality bases, and merging paired reads (--collapse), using the additional parameters: --adapter1 AGATCGGAAGAGCACACGTCTGAACTCCAGTCAC --adapter2 AGATCGGAAGAGCGTCGTGTAGGGAAAGAGTGT --trimns --trimqualities --minlength 30 --minadapteroverlap 1.

Initial taxonomic composition and contamination were assessed with FastQ Screen v0.15.2 (8). Collapsed reads were aligned with Bowtie2 v2.4.4 (9) against multiple reference genomes (goat, cattle, human, horse, sheep, pig, dog, deer, fox). Collapsed reads were aligned to the goat reference genome ARS1 (10) with BWA aln v0.7.17-r1188 (11), using parameters tailored for ancient DNA (-l 1024 -n 0.01 -o 2) (12). Generated .sai files were converted to .sam (bwa samse) with read group (@RG) added, then to BAM with Samtools v1.13 (13). BAM files were processed in the following steps: sorted (samtools sort), filtered to retain reads with mapping quality ≥30 (samtools view -q 30), and duplicates removed (samtools rmdup -s).

As competitive alignment using Fastq Screen identified both human and fox-associated reads, we aligned collapsed reads to both the human (hg38) and fox (vulvul2-2) genomes, using bwa aln with the relaxed parameters described above, without any additional read filtering. Reads which aligned to either human or fox genomes were then removed from the goat ARS1 bam files. Final quality checks on bam files were done with samtools flagstat, samtools stats, and Qualimap v2.2.1 (14). MapDamage (15) was applied to assess ancient DNA damage patterns and reads were softclipped for 6 bp on either end.

#### Genetic analyses

##### LASER

A projection Principal Component Analysis (PCA) was performed using LASER v2.4 (16) to maximize the inclusion of the low-coverage Murciélagos1 sample. The PCA reference space and projection transformation were constructed using the VarGoats dataset (17, 18), where individuals with high relatedness removed as in (19), filtered to retain transversions with a MAF of 5%, resulting in a total of 2,420,382 SNPs. All ancient samples were then projected onto the PCA space, and then filtered for individuals covered by less than 5,000 loci. To reduce stochastic variation and provide robust estimates, 100 replicates were conducted, enabling the calculation of standard deviations.

##### Pseudohaploid genotypes

To further assess the genetic affinities of the Murciélagos goat, we generated pseudohaploid genotypes using ANGSD v0.941-11-g7a5e0db (20), applying the -doHaploCall function to randomly sample one high-quality base per position. We applied ancient DNA-specific filters (-uniqueOnly 1, -remove_bads 1, -trim 4, -minMapQ 30, -minQ 25, -rmTrans 1) and restricted the analysis to 8,026,660 autosomal transversion SNPs from the VarGoats dataset (19). Genotypes were converted to PLINK format using haploToPlink, merged with imputed modern genotypes from VarGoats, and converted to EIGENSTRAT format using convertf from the ADMIXTOOLS suite (21).

##### Outgroup $f_3$

To estimate the degree of shared genetic drift between Murciélagos1 and modern breeds, outgroup $f_3$ statistics (22) were calculated using ADMIXTOOLS version 7.0.2 (21). We used granular breed groupings where possible and domestic sheep were used as the outgroup.

##### *D* statistics

To test for evidence of gene flow and estimate its extent between Murciélagos1, Neolithic Serbia (the Blagotin group), and Neolithic Eastern Iran (Semnan3), and modern breeds, *D* statistics were calculated using ADMIXTOOLS version 7.0.2 (21). The model to test for wild or domestic status was structured as *D*(Outgroup, Neolithic; *Capra pyrenaica*, Modern breeds). The model to test for genetic differentiation between Neolithic goats and modern breeds was structured as *D*(Outgroup, Neolithic; Bermeya (Modern North Spain), Modern breeds). To test if Murciélagos1 shared more ancestry with Neolithic Serbian goats (Blagotin) or modern breeds *D* statistics were calculated as *D*(Outgroup, Murciélagos1: Blagotin, Modern breeds). The Blagotin goats were grouped and considered as one population.

<u>Mitochondrial DNA</u>

To complement nuclear data, mitochondrial DNA from the Murciélagos1 goat sample was analyzed. Collapsed reads were aligned separately to a circularized version of the goat mitochondrial genome (15 bp duplicated at both ends) using bwa aln v0.7.17 (11) with ancient DNA-specific parameters (-l 1024 -n 0.01 -o 2). BAM files were filtered to retain only reads ≥30 bp, with mapping quality ≥30, sorted, and deduplicated using Samtools v1.13 (13) and Picard tools. To exclude potential contaminants, reads were also aligned against human (GRCh38) and red fox (VulVul2.2) mitochondrial genomes using the same pipeline as before and removed from the goat mitochondrial bam file.

To obtain a representative mitochondrial sequence for downstream analyses, consensus mitochondrial sequences were generated from filtered BAM files (human and fox contaminants removed) using ANGSD v0.941 (20) with the -doFasta 2 option to call consensus bases, applying minimum depth 5, base quality ≥20, and mapping quality ≥30. The output fasta sequences were subsequently soft-clipped by removing 15 bp from both 5′ and 3′ ends to correct for circular genome overlap. This step was performed with a custom Python script that concatenated sequence lines before trimming and rewriting the fasta file.

We compiled and aligned the Murciélagos1 mitochondrial sequence along with published modern and ancient goat mitochondrial genomes, and an outgroup sheep sequence. The sequences were merged into a single FASTA file and aligned using MAFFT v7 (23) with default parameters. This multiple sequence alignment was used for subsequent phylogenetic analyses. Phylogenetic trees were inferred using RAx mL v8.2.12 (24) under the GTR+Gamma substitution model. We performed 100 rapid bootstrap replicates combined with a thorough maximum likelihood search using the '-f a' option to estimate node support. Phylogenetic trees were visualized and edited with FigTree v1.4.4 (http://tree.bio.ed.ac.uk/software/figtree/)to improve clarity and presentation. Branch lengths, bootstrap values, and clade labels were manually adjusted.

<u>Metagenomics</u>

In order to characterize the microbiome of the leather sample, fastq files were cleaned from host reads by aligning libraries with Bowtie2 v2.4.4 (9) on a concatenated fasta mix of different animal and human reference genomes (GRCh38, Sscrofa11.1, ARS-UCD1.2, mCerEla1.1, ASM170441v1, EquCab3.0, ARS-UI_Ramb_v2.0) with no mapping quality filter applied. Then, the metagenomic profile of the non-aligned reads was computed with KrakenUniq v1.0.4 (25) using a custom database containing a microbial version of NCBI nt combined with human and complete eukaryotic reference genomes (26).

<u>Radiocarbon dating</u>

The leather was dated by the AMS method at the Tandem Laboratory, Uppsala University. Lipids were extracted from a subsample of the leather using a mixture of ether and ethanol (3:1, w/w) for 1 hour at 40 °C. The sample was then rinsed with distilled water, after which 1% HCl was added and the mixture was maintained for 4 hours at 40°C. Following this, the sample was rinsed to neutrality with distilled water and subsequently treated with 0.25% NaOH for 45 minutes at room temperature. The insoluble fraction (A) was rinsed again to neutrality, acidified to pH 3 using 1% HCl, and finally dried at 40°C. Prior to accelerator mass spectrometry (AMS) determination of the $^{14}C$ content, the material was combusted to $CO_2$ and graphitized using an Fe-catalyst reaction. In the present investigation, the insoluble fraction (INS) was dated. This date (Ua-78252) has been compared with other available radiocarbon dates from the Cueva de los Murciélagos on short-lived terrestrial samples and calibrated using OxCal v.4.4.4. A calibrated model was developed using OxCal command *Phase* and calibration curve *IntCal20* to gather the values for the ranges of the cave use (27, 28).

**Supplementary Note: Sexing**
Sex determination of Murciélagos1 was performed using two complementary molecular approaches, to increase the reliability of the inference. The first method, based on Park et al. (2015)(29), uses a residual-based approach that exploits the linear relationship between chromosome length and sequencing read depth. In males, the X chromosome is present in a single copy (haploid), causing its read depth to deviate from this linear trend.

Chromosome-level read counts were obtained, and residuals were calculated to infer sex (Fig. S7). According to this method, Murciélagos1 appears to be female, but falls at the lower end of the female range, which could be due to the highly fragmented nature of the ancient DNA.

The second method involves calculating the ratio of sequencing reads mapped to the Y chromosome versus the X chromosome (Y:X ratio). Because females lack a Y chromosome, this ratio should be close to zero for females. Reads aligning to known sex chromosome scaffolds were counted and the ratio computed. This approach also suggests that Murciélagos1 is female, though the individual again lies at the lower limit of the female range (Fig. S8). Given the consistent but borderline results from both methods, these findings should be interpreted with caution

**Supplementary Note: Human DNA alignments**
After removing reads aligning to goat and fox in UDG and non-UDG treated libraries, the remaining library reads were mapped against the Genome Reference Consortium Human Build 37 (GRCH37 or hg19) and the human mitochondrial revised Cambridge Reference Sequence (rCRS) (30) with nf-core/eager (31). Endogenous human DNA content in the individual libraries ranges between 0.14-0.18% post-deduplication. Deamination in terminal 5' and 3' positions oscillates between 1.6 and 5.8% in UDG-treated libraries and between 8.3 and 15.1%, lower than in the goat reads in the same libraries, but higher than in the fox reads (Fig. S5 and Table S1). Mean fragment size, between 66 and 77bp, is also higher than that reported for the goat reads in the same libraries (Table S1, Fig. 3,S4).

Libraries were merged post-deduplication and the resulting merged reads aligned again against hg19 and rCRS with nf-core EAGER. Ten terminal base pairs were removed from the non-UDG treated libraries prior to genotyping. Whole-genome genotyping was conducted using the pileupCaller and the option "randomHaploid", with genotypes being called at the Ancient Human DNA Target Enrichment panel (Twist Bioscience) nucleotide site list. Circular mapper was used for mtDNA genotyping against rCRS (31). Only 7,261 SNPs from the 1,372K targeted in the Twist panel (0.53%) could be identified and just 133 reads could be aligned against the human mtDNA reference, therefore insufficient for broad ancestry analysis.

Sex determination was also attempted in the merged human reads using the DetERRmine script (https://github.com/TCLamnidis/Sex.DetERRmine, (32)) built in the nf-core/EAGER pipeline. This method estimates chromosomal sex by calculating the coverage of the mapped reads on the X and Y chromosomes relative to the one on the autosomes. Biological females are expected to have an X-rate of 1, as they have the same number of X chromosomes as autosomes, and a Y-rate of 0. In turn, biological males are expected to have both X and Y-rates of 0.5. Ratios of human reads aligning on the X and Y chromosomes versus autosomes in the merged library are, respectively, 0.5 and 0.4, thus compatible with a male specimen (Table S1, Fig. S11). This result should be, however, interpreted with extreme caution, as the number of reads mapping against the X and Y chromosomes was very low - 154 and 201, respectively - representing approximately 2% of the 7,261 reads covering SNP positions from the Twist panel (Table S1).

**Supplementary figures**

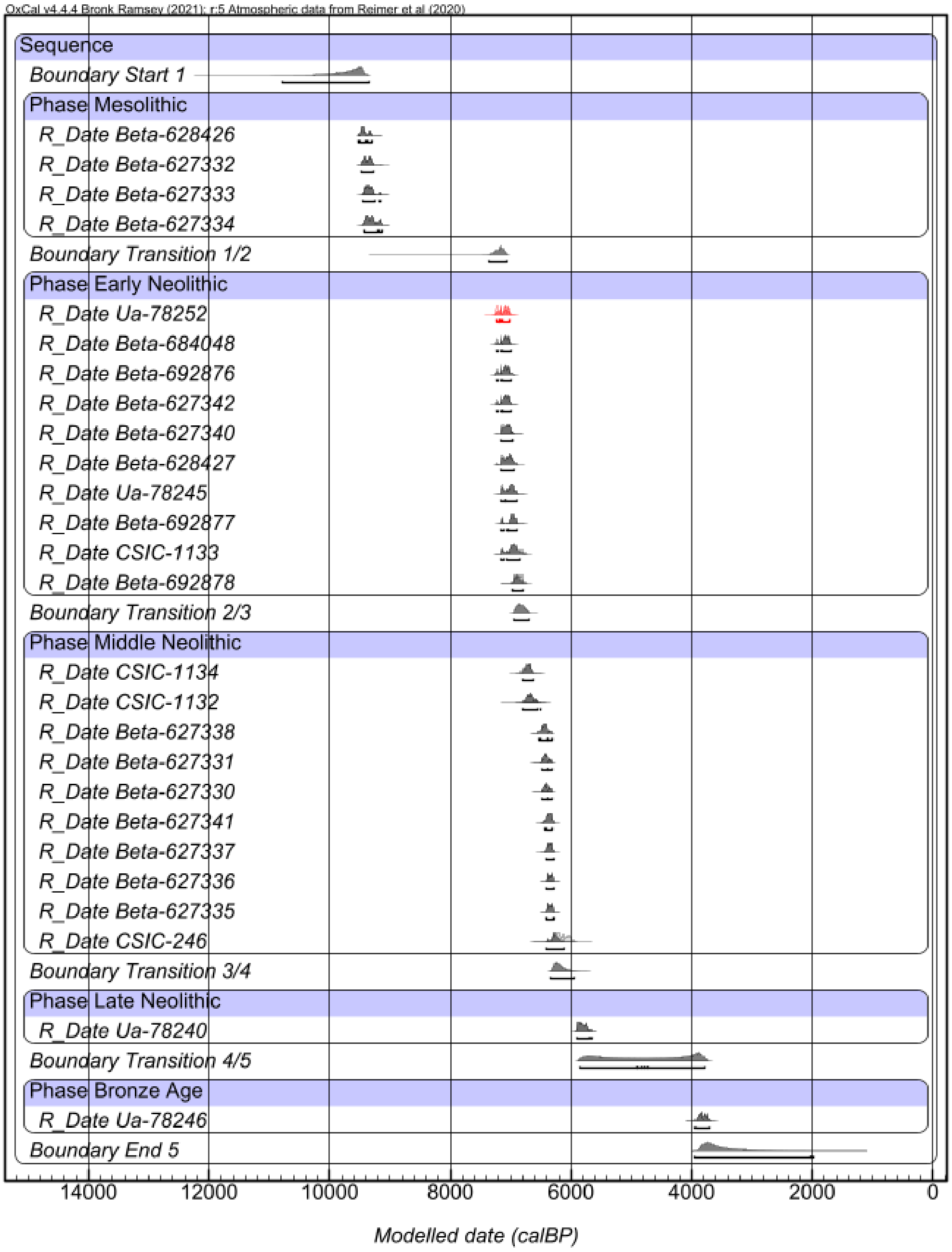


Figure S1: Probability distribution of previously published radiocarbon dates from Cueva de los Murciélagos, including the date of the leather from this study (in red). Sequence chronological ranges for the estimated start and end of each phase and the modelled ranges of each radiocarbon date were calculated using OxCal v4.4.4.

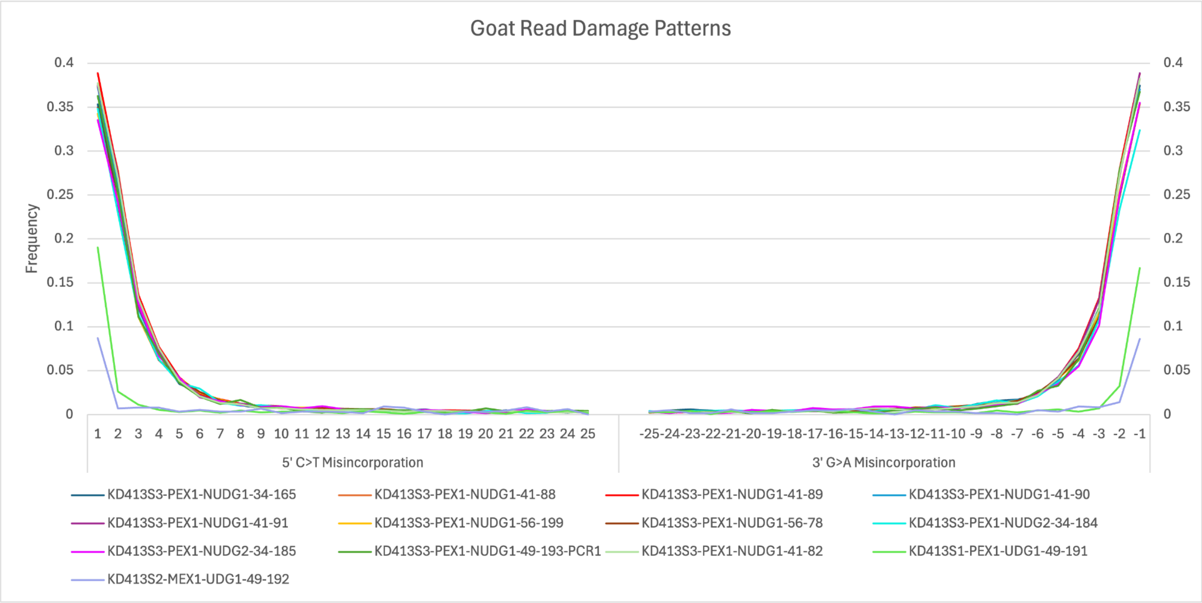


Figure S2: MapDamage results showing the characteristic ancient DNA mutations on reads aligned to the goat genome after filtering out fox and human reads.

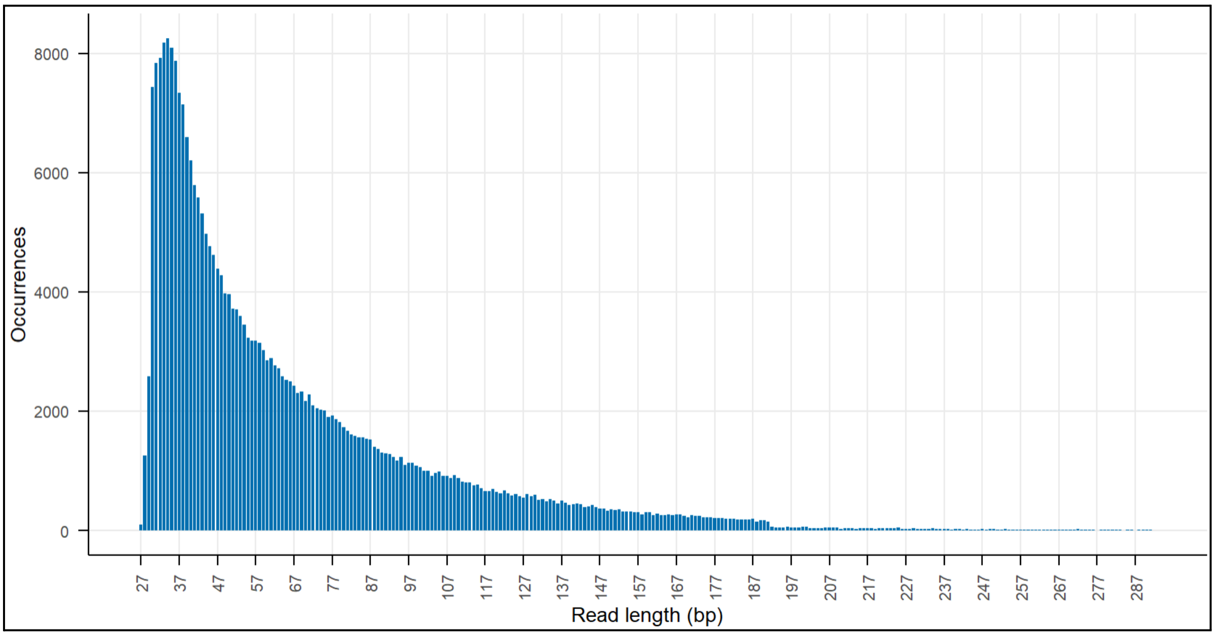


Figure S3: MapDamage results showing the read length distribution (bp) of reads aligned to the human genome after filtering out fox and goat reads.

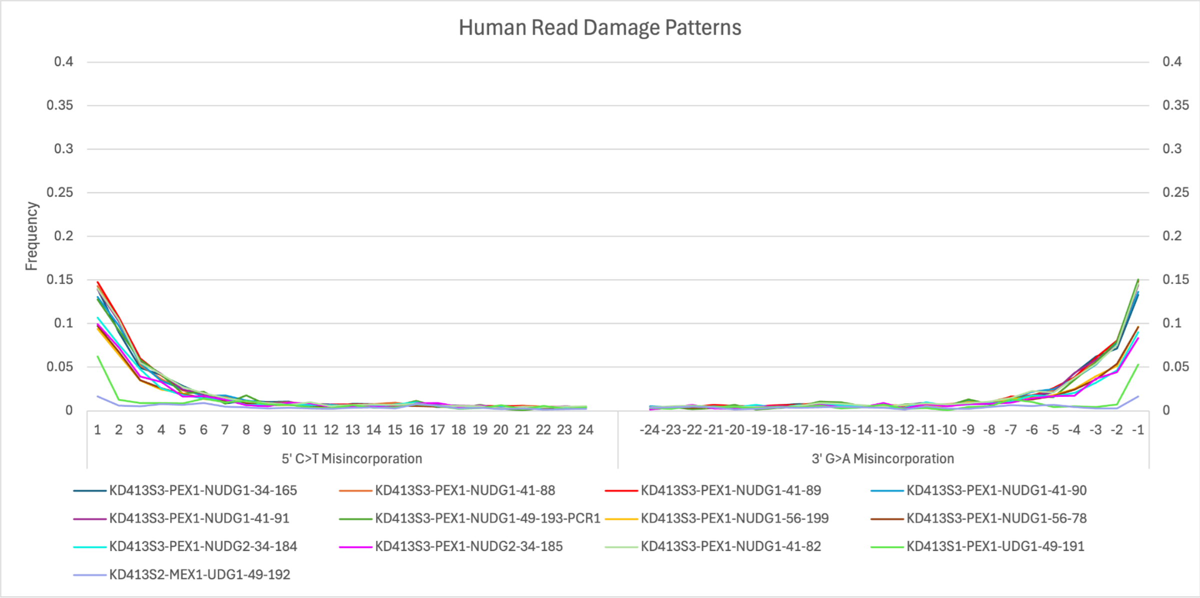


Figure S4: MapDamage results showing the characteristic ancient DNA mutations on reads aligned to the human genome after filtering out fox and goat reads.

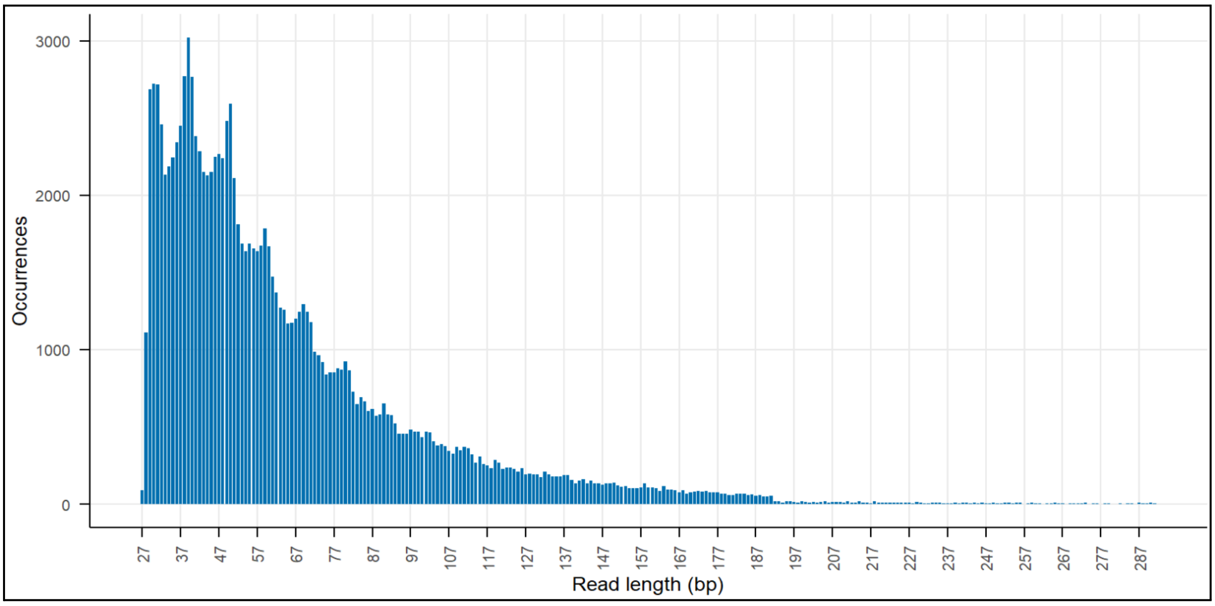


Figure S5: MapDamage results showing the read length distribution (bp), on reads aligned to the fox genome after filtering out human and goat reads.

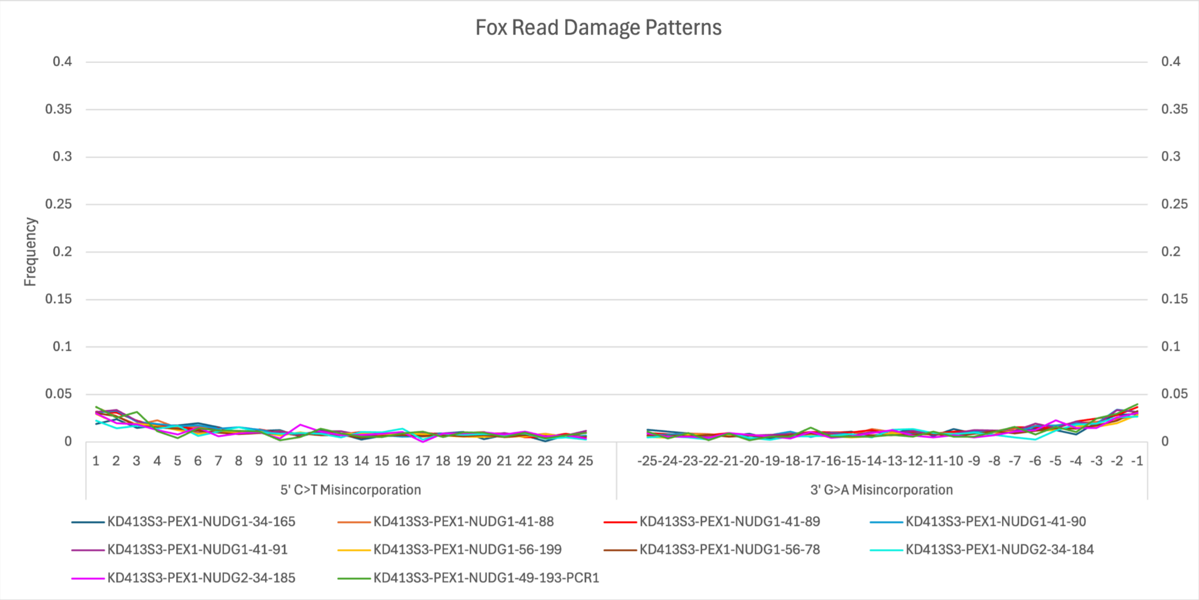


Figure S6: MapDamage results showing the characteristic mutations of ancient DNA on reads aligned to the fox genome after filtering out human and goat reads.

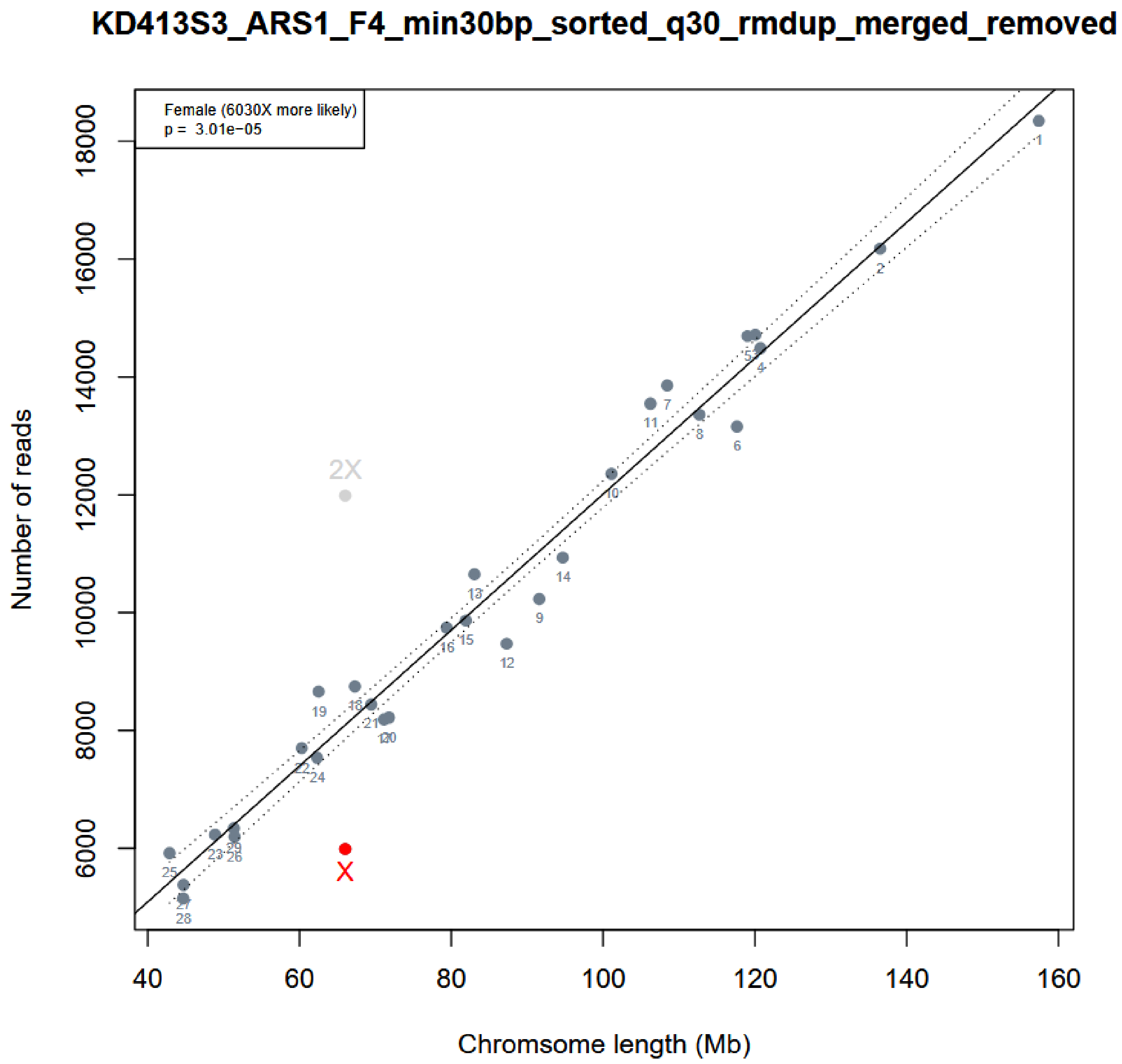


Figure S7: Sex determination of the Murciélagos1 goat (shown in red), using the residual-based method following Park et al. (2015)(29). Gray samples are test samples.

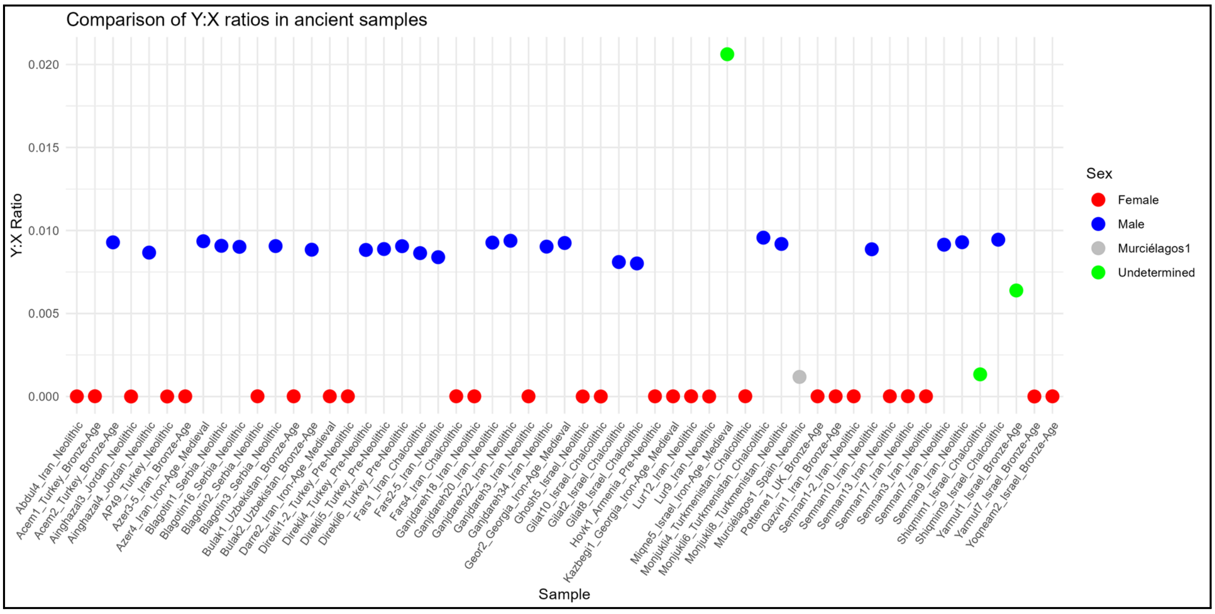


Figure S8: Sex determination of the Murciélagos1 goat: based on the X:Y chromosome read ratio. Red indicates female, blue indicates male, green indicates undetermined, and grey indicates Murciélagos1.

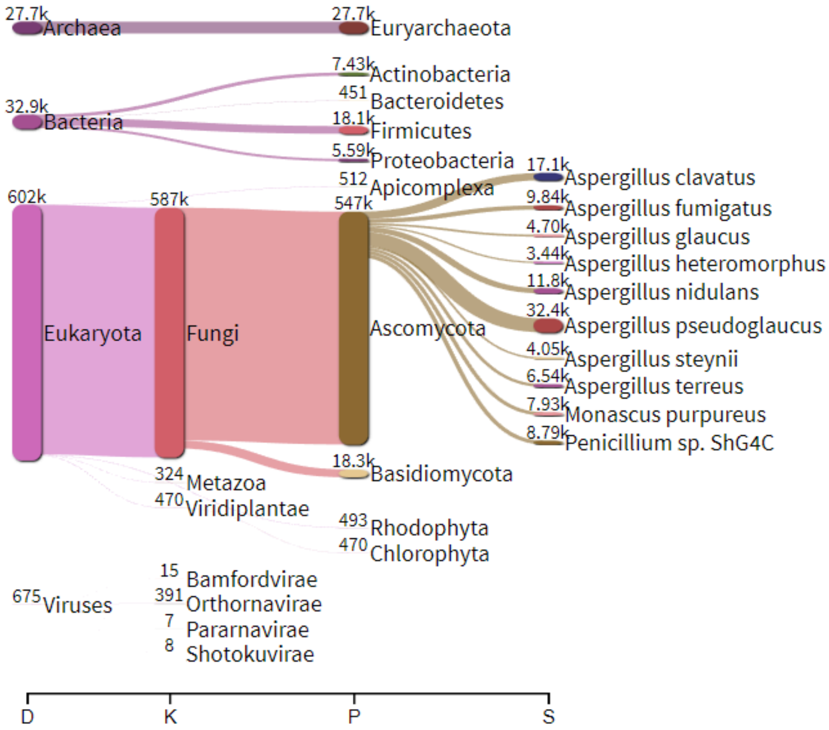


Figure S9: Taxonomic composition of the metagenomic reads recovered from the leather Murciélagos1. Sankey diagram showing the taxonomic classification of non-host metagenomic reads assigned at different levels (Domain [D], Kingdom [K], Phylum [P], and Species [S]). Most reads are assigned to the *Eukaryota* domain, particularly to *Fungi*

(*Ascomycota*), with dominant genera such as *Aspergillus* and *Penicillium*. Bacterial and viral taxa are also detected in lower abundance.

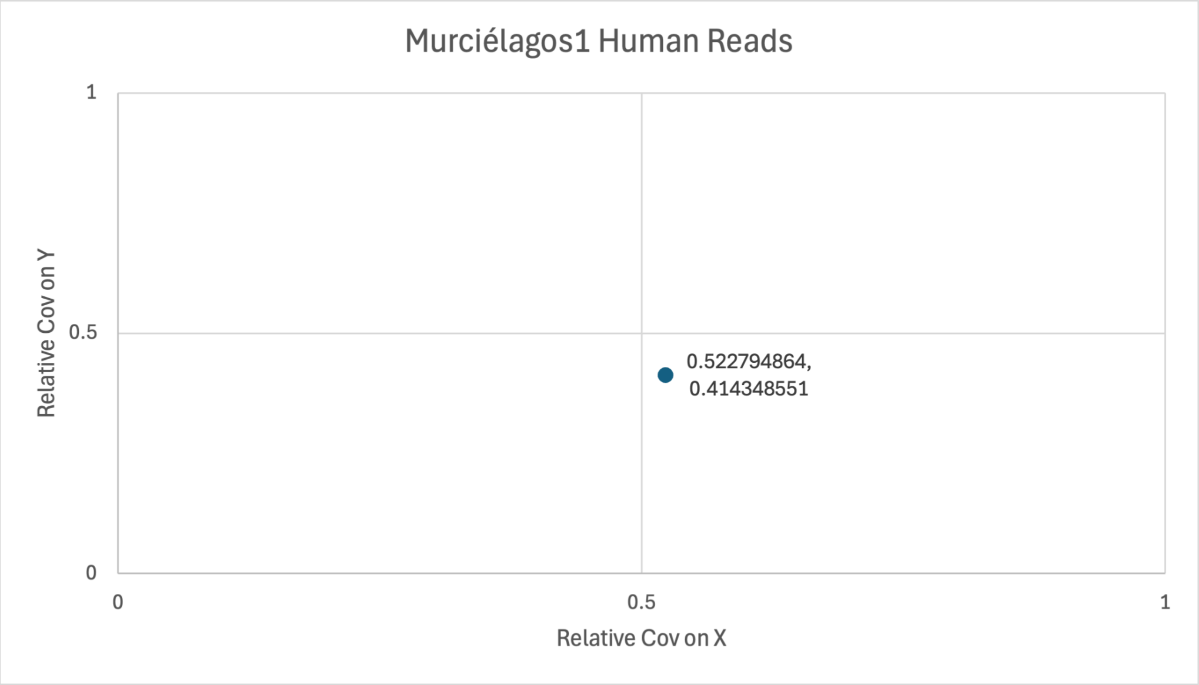


Figure S10: Sex determination of human reads from the Murciélagos1 sample using the Sex.DetERRmine script and the Ancient Human DNA Target Enrichment panel (Twist Bioscience) nucleotide site list bedfile (32).

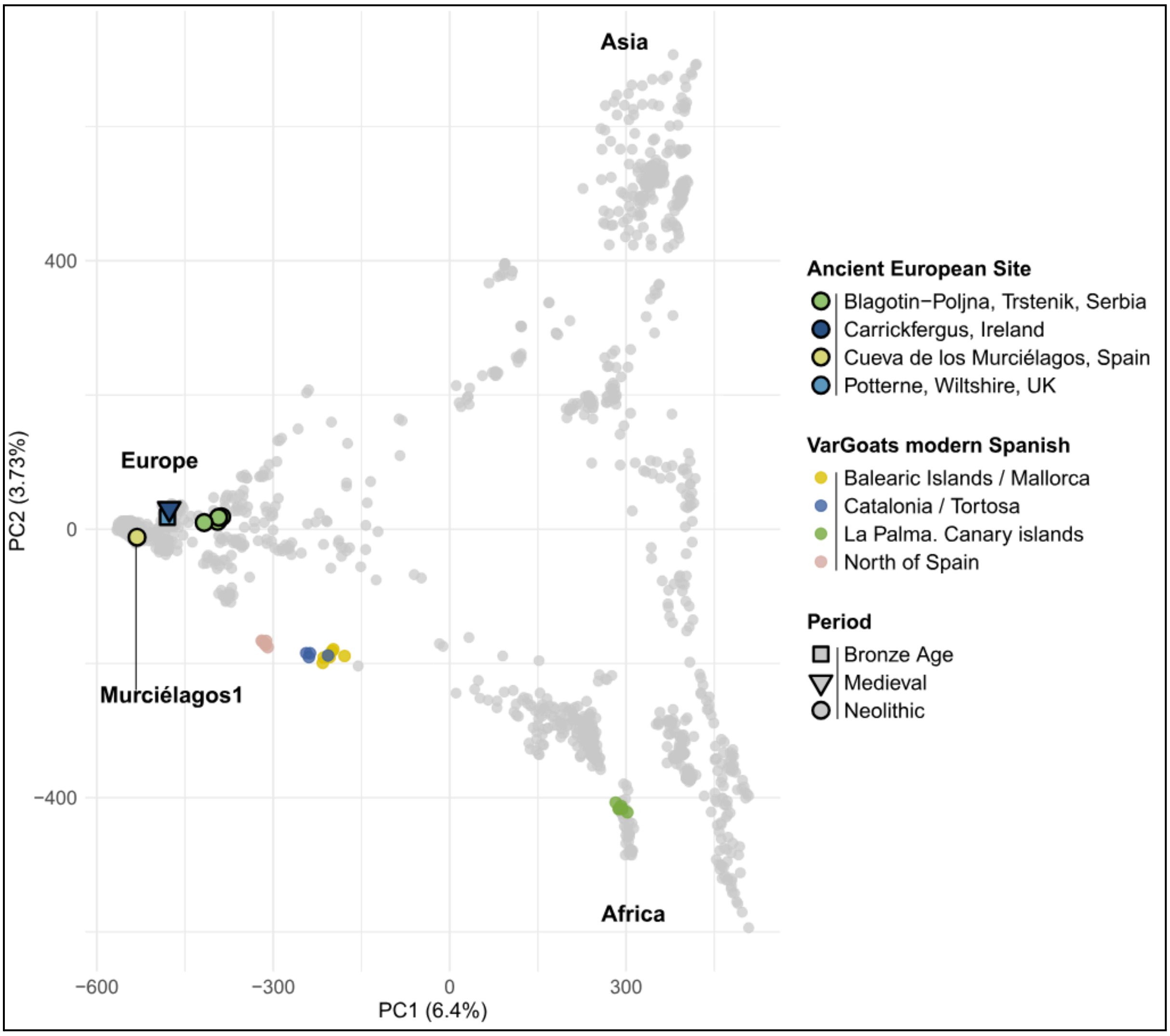


Figure S11: Principal component analysis (PCA) of the VarGoats Modern dataset (54) and ancient European goats, including our study sample Murciélagos1. Each point represents the genetic information of a goat sample. Axes PC1 (6.4%) and PC2 (3.73%) represent the first two principal components, which explain the genetic variation. The period is indicated by the shape of the point, and ancient samples have a black border. The sampling location is indicated by the color of the point. Grey points correspond to modern non-European samples; those located at the bottom right of the PCA form the African cluster, and those at the top right form the Asian cluster.

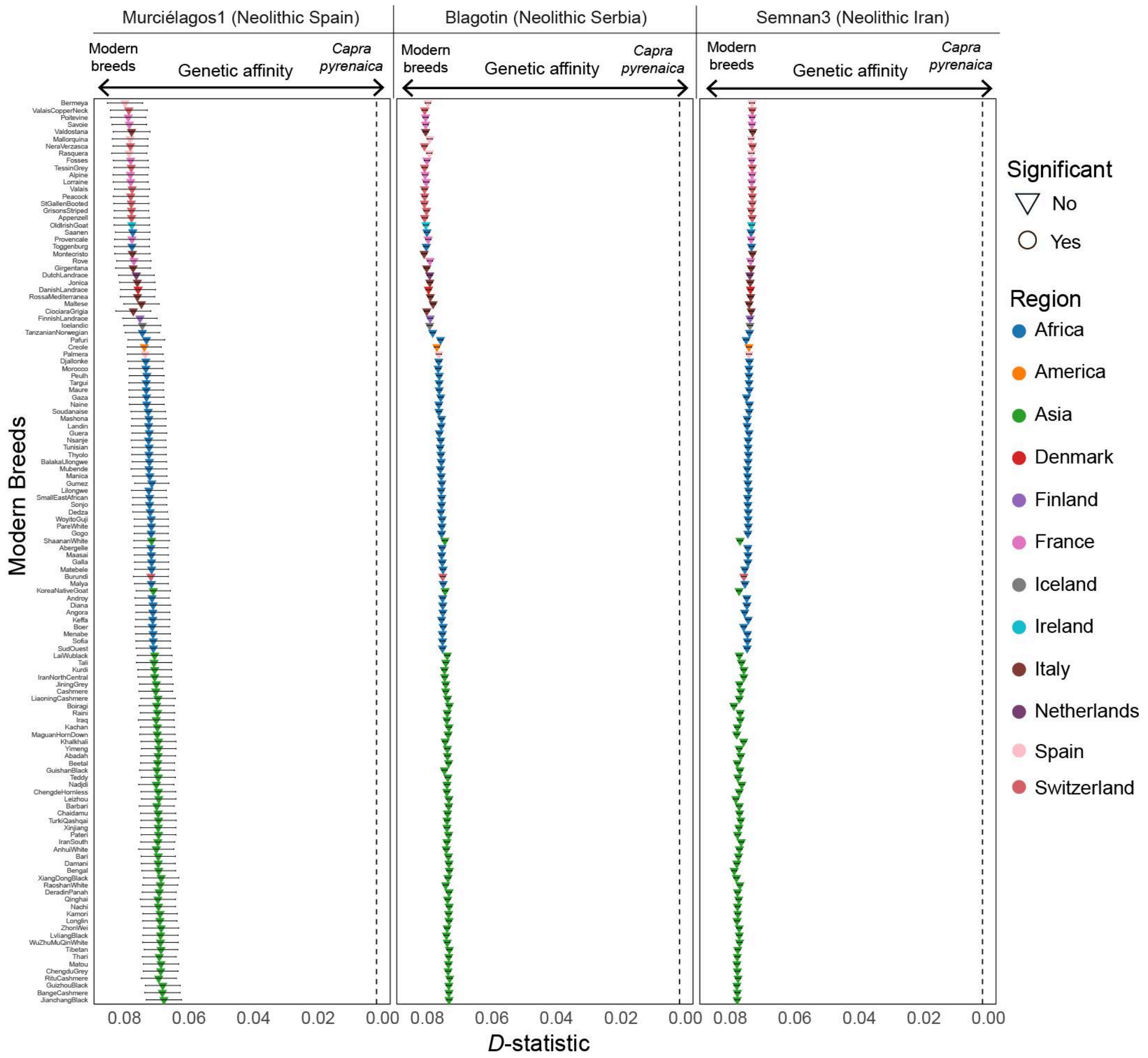


Figure S12: *D* statistics of the form *D*(Out, Neolithic; *Capra pyrenaica*, Modern breeds), where Neolithic was varied between Murciélagos1, Neolithic Serbian goat (grouped together as Blagotin), and a Neolithic east Iranian goat (Semnan3), the later two living roughly a millennia within Murciélagos1 estimated age. This *D* test assesses the affinity of Murciélagos1, Blagotin and Semnan with The Iberian ibex (*Capra pyrenaica)* compared with modern breeds. Negative indicates closer affinity (more shared derived alleles) to *Capra pyrenaica* and positive indicates closer affinity towards the modern breed in question (Y axis).

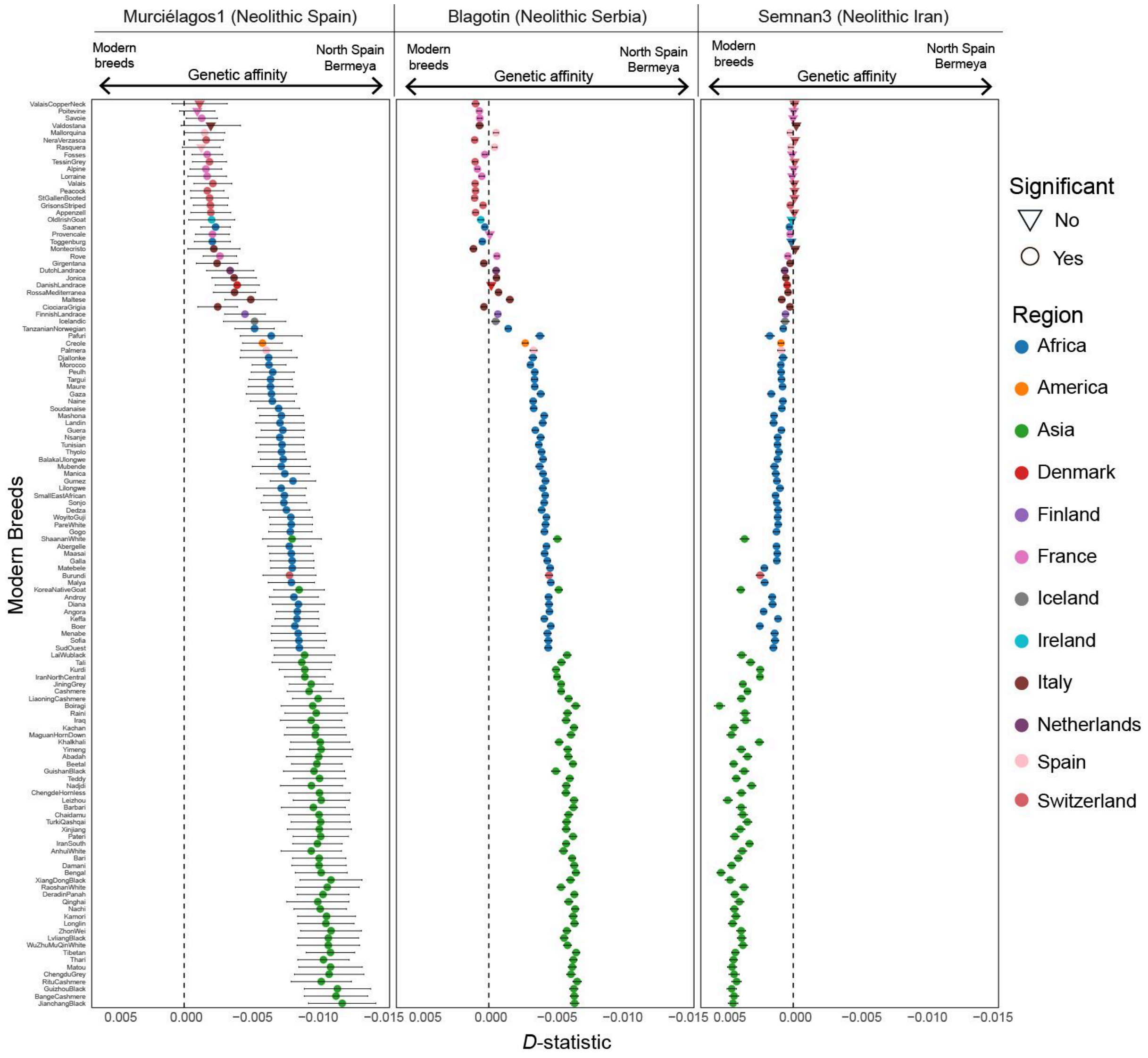


Figure S13: *D* statistics of the form *D*(Out, Neolithic; Bermeya (Modern northern Spain), Modern breeds), where Neolithic was varied between Murciélagos1, Neolithic Serbian goat (grouped together as Blagotin), and a Neolithic east Iranian goat (Semnan3), the later two living roughly a millennia within Murciélagos1 estimated age. This *D* test assesses the affinity of Murciélagos1, Blagotin and Semnan with Bermeya compared with modern breeds. Negative indicates closer affinity (more shared derived alleles) to Bermeya and positive indicates closer affinity towards the modern breed in question (Y axis).

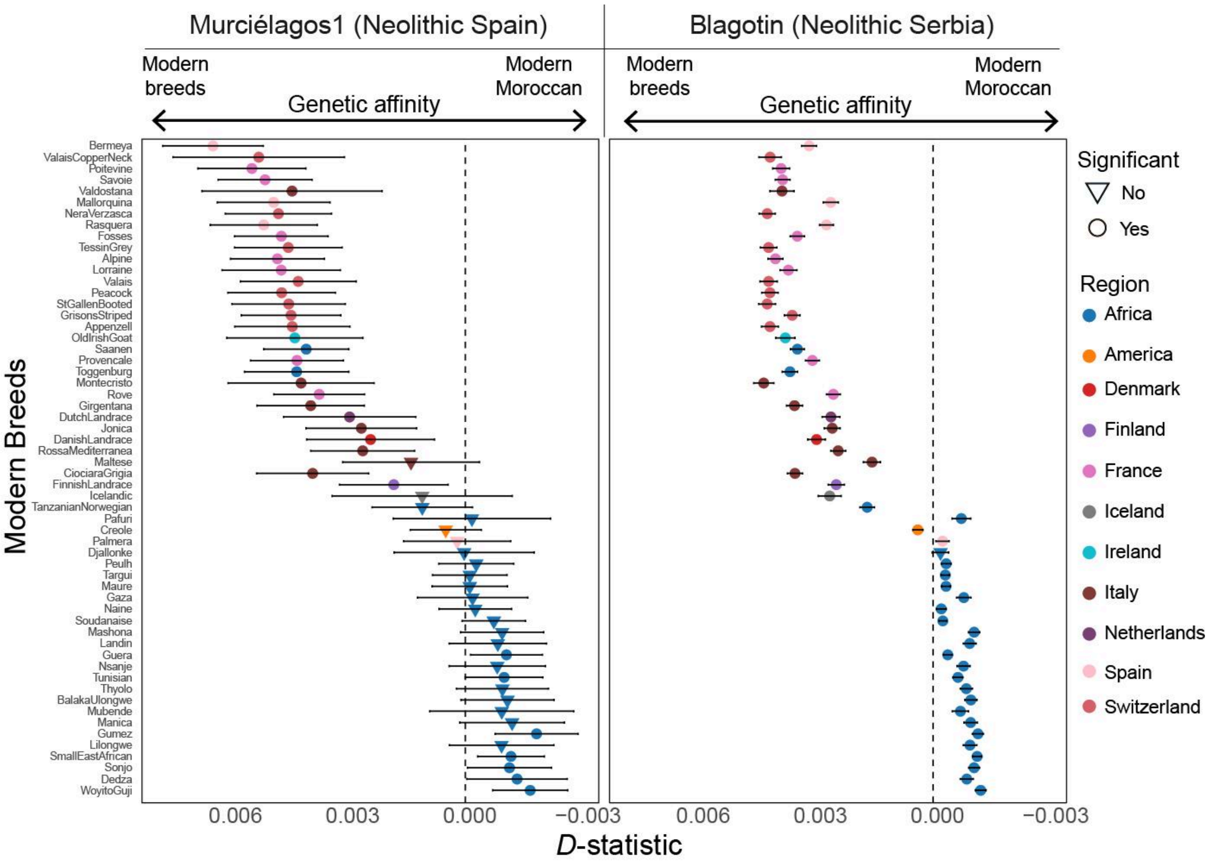


Figure S14: *D* statistics of the form *D*(Out, Neolithic; Moroccan, European/African Modern breeds), where Neolithic was varied between Murciélagos1, and Neolithic Serbian goat (grouped together as Blagotin), living roughly a millennia within Murciélagos1 estimated age. This *D* test assesses the affinity of Murciélagos1 and Blagotin with Moroccan goats compared with European/African modern breeds. Negative indicates closer affinity (more shared derived alleles) to Moroccan goats and positive indicates closer affinity towards the modern breed in question (Y axis).

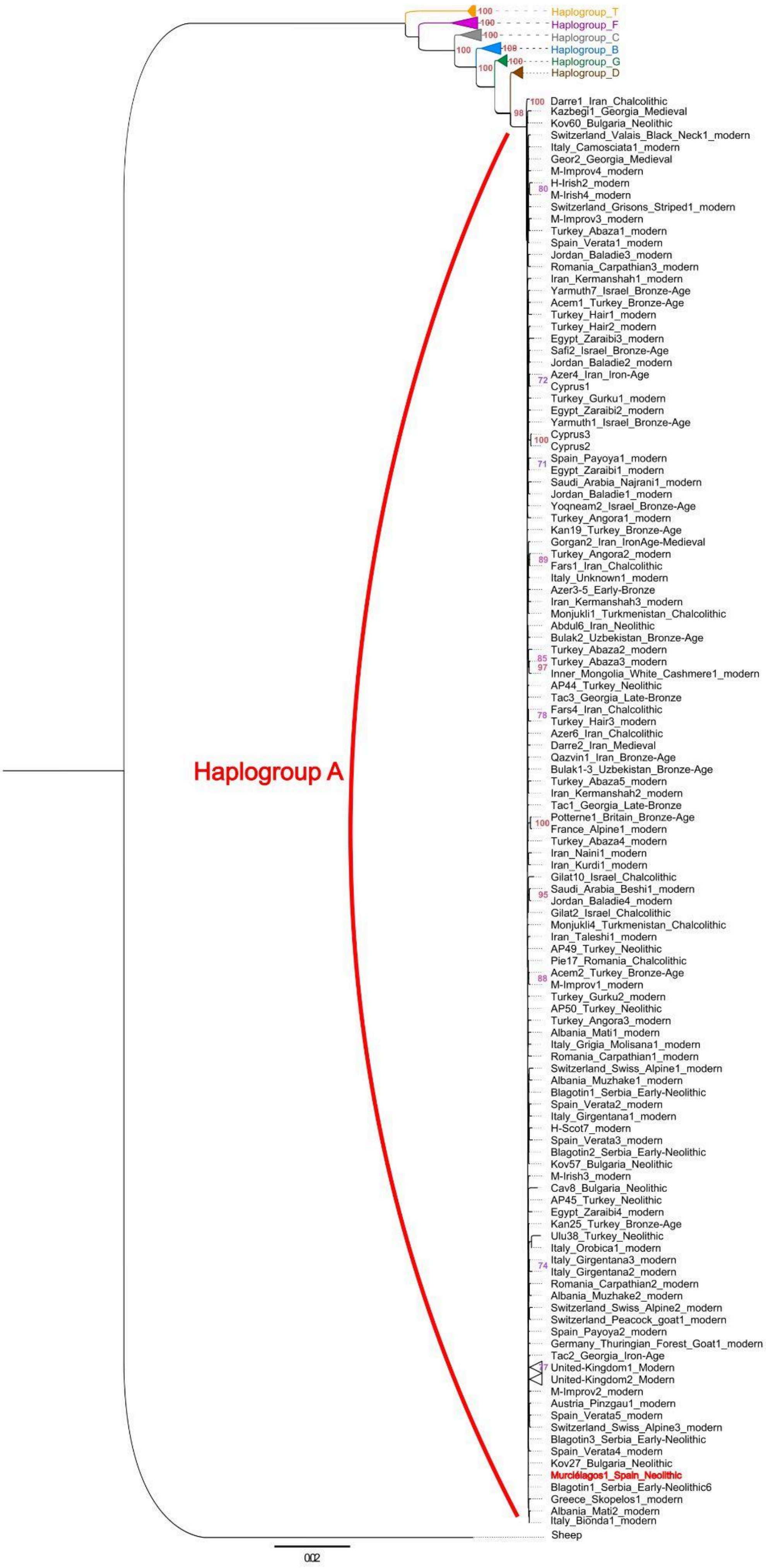

Haplogroup_T
Haplogroup_F
Haplogroup_C
Haplogroup_B
Haplogroup_G
Haplogroup_D
Darre1_Iran_Chalcolithic
Kazbegi1_Georgia_Medieval
Kov60_Bulgaria_Neolithic
Switzerland_Valais_Black_Neck1_modern
Italy_Camosciata1_modern
Geor2_Georgia_Medieval
M-Improv4_modern
H-Irish2_modern
M-Irish4_modern
Switzerland_Grisons_Striped1_modern
M-Improv3_modern
Turkey_Abaza1_modern
Spain_Verata1_modern
Jordan_Baladie3_modern
Romania_Carpathian3_modern
Iran_Kermanshah1_modern
Yarmuth7_Israel_Bronze-Age
Acem1_Turkey_Bronze-Age
Turkey_Hair1_modern
Turkey_Hair2_modern
Egypt_Zaraibi3_modern
Safi2_Israel_Bronze-Age
Jordan_Baladie2_modern
Azer4_Iran_Iron-Age
Cyprus1
Turkey_Gurku1_modern
Egypt_Zaraibi2_modern
Yarmuth1_Israel_Bronze-Age
Cyprus3
Cyprus2
Spain_Payoya1_modern
Egypt_Zaraibi1_modern
Saudi_Arabia_Najrani1_modern
Jordan_Baladie1_modern
Yoqneam2_Israel_Bronze-Age
Turkey_Angora1_modern
Kan19_Turkey_Bronze-Age
Gorgan2_Iran_IronAge-Medieval
Turkey_Angora2_modern
Fars1_Iran_Chalcolithic
Italy_Unknown1_modern
Azer3-5_Early-Bronze
Iran_Kermanshah3_modern
Monjukli1_Turkmenistan_Chalcolithic
Abdul6_Iran_Neolithic
Bulak2_Uzbekistan_Bronze-Age
Turkey_Abaza2_modern
Turkey_Abaza3_modern
Inner_Mongolia_White_Cashmere1_modern
AP44_Turkey_Neolithic
Tac3_Georgia_Late-Bronze
Fars4_Iran_Chalcolithic
Turkey_Hair3_modern
Azer6_Iran_Chalcolithic
Darre2_Iran_Medieval
Qazvin1_Iran_Bronze-Age
Bulak1-3_Uzbekistan_Bronze-Age
Turkey_Abaza5_modern
Iran_Kermanshah2_modern
Tac1_Georgia_Late-Bronze
Potterne1_Britain_Bronze-Age
France_Alpine1_modern
Turkey_Abaza4_modern
Iran_Naini1_modern
Iran_Kurdi1_modern
Gilat10_Israel_Chalcolithic
Saudi_Arabia_Beshi1_modern
Jordan_Baladie4_modern
Gilat2_Israel_Chalcolithic
Monjukli4_Turkmenistan_Chalcolithic
Iran_Taleshi1_modern
AP49_Turkey_Neolithic
Pie17_Romania_Chalcolithic
Acem2_Turkey_Bronze-Age
M-Improv1_modern
Turkey_Gurku2_modern
AP50_Turkey_Neolithic
Turkey_Angora3_modern
Albania_Mati1_modern
Italy_Grigia_Molisana1_modern
Romania_Carpathian1_modern
Switzerland_Swiss_Alpine1_modern
Albania_Muzhake1_modern
Blagotin1_Serbia_Early-Neolithic
Spain_Verata2_modern
Italy_Girgentana1_modern
H-Scot7_modern
Spain_Verata3_modern
Blagotin2_Serbia_Early-Neolithic
Kov57_Bulgaria_Neolithic
M-Irish3_modern
Cav8_Bulgaria_Neolithic
AP45_Turkey_Neolithic
Egypt_Zaraibi4_modern
Kan25_Turkey_Bronze-Age
Ulu38_Turkey_Neolithic
Italy_Orobica1_modern
Italy_Girgentana3_modern
Italy_Girgentana2_modern
Romania_Carpathian2_modern
Albania_Muzhake2_modern
Switzerland_Swiss_Alpine2_modern
Switzerland_Peacock_goat1_modern
Spain_Payoya2_modern
Germany_Thuringian_Forest_Goat1_modern
Tac2_Georgia_Iron-Age
United-Kingdom1_Modern
United-Kingdom2_Modern
M-Improv2_modern
Austria_Pinzgau1_modern
Spain_Verata5_modern
Switzerland_Swiss_Alpine3_modern
Blagotin3_Serbia_Early-Neolithic
Spain_Verata4_modern
Kov27_Bulgaria_Neolithic
Murciélagos1_Spain_Neolithic
Blagotin1_Serbia_Early-Neolithic6
Greece_Skopelos1_modern
Albania_Mati2_modern
Italy_Bionda1_modern
Sheep
Haplogroup A
0.02

Figure S15 : Maximum Likelihood phylogenetic tree based on mitochondrial DNA sequences. The tree was constructed using RAxML with the GTR+Gamma substitution model and 100 bootstrap replicates. The dataset includes ancient goat samples, with Murciélagos1 highlighted in red. Sheep (Ovis aries) sequences were included as an outgroup to root the tree. Samples belonging to the same haplogroup, other than A, were collapsed into single representative branches for clarity.

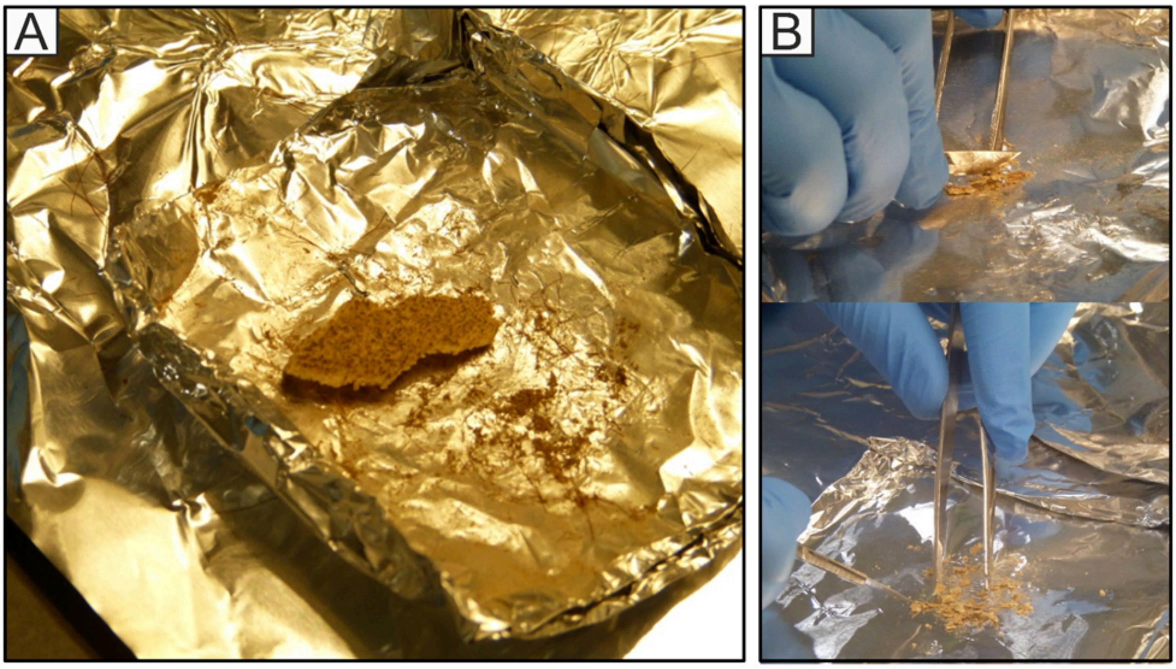


Figure S16: A) Murciélagos 1 (P-082); B) Processing leather in the Smurfit institute of Genetics laboratory, Trinity College Dublin.
